# Gene Duplication, Horizontal Gene Transfer, and Trait Trade-offs Drive Evolution of Post-Fire Resource Acquisition in Pyrophilous Fungi

**DOI:** 10.1101/2025.07.17.665399

**Authors:** Ehsan Sari, Dylan J. Enright, Maria Ordonez, Steven D. Allison, Peter M. Homyak, Michael J. Wilkins, Sydney I. Glassman

## Abstract

Wildfires significantly alter soil carbon (C) and nitrogen (N), reducing microbial richness and biomass, while selecting for “fire-loving” pyrophilous microbes that drive post-fire nutrient cycling. However, the genomic strategies and functional trade-offs (balancing gains in one trait with costs in another) underlying the traits that enable pyrophilous microbes to survive and thrive post-fire are virtually unknown. We hypothesized that pyrophilous fungi employ specialized genomic adaptations for C and N cycling, with evolutionary trade-offs between traits governing aromatic C degradation, N acquisition pathways, and rapid growth. To test these hypotheses, we performed complementary comparative genomics, transcriptomics after pyrogenic organic matter amendment, and growth rate bioassays for 18 pyrophilous fungi from five Ascomycota (Eurotiales, Pleosporales, Sordariales, Coniochaetales, and Pezizales) and three Basidiomycota (Agaricales, Holtermaniales, and Geminibasidiales) orders isolated from burned soils. We found a dramatic trait trade-off between fast growth and number of genes responsible for aromatic C degradation, implying burned environments select for metabolically costly genes despite their evolutionary cost. We used the comparative genomics framework to evaluate genomic signatures of evolution and found that either gene duplication and somatic mutation, or recombination via sexual reproduction, were the primary drivers of fungal genomic variation in aromatic C degradation and N acquisition genes. Finally, we identified cross-kingdom bacterial to fungal horizontal gene transfer as a secondary strategy producing novel aromatic C degradation genes. Overall, we found that trait trade-offs and genome evolutionary strategies are key drivers that may predict the persistence and contribution of pyrophilous fungi to global C and N cycling.

**Significance statement:** Wildfires affect large tracts of the terrestrial biosphere, and while much is understood about plant adaptations to fire, here we uncovered for the first time genomic adaptations in fungi that can be modeled to predict impacts on global C and N cycling. We tested theorized trait trade-offs, and showed with genomics and bioassays how evolutionary trade-offs, like prioritizing aromatic C degradation at the expense of rapid growth, enable pyrophilous fungi to thrive post-fire. Further, we advance fungal genomics, and reveal the importance of mechanisms like gene duplication, sexual reproduction, and cross-kingdom horizontal gene transfer for enabling adaptation and evolutionary diversification in fungi. Finally, we identify fungi that may be useful inoculants to enhance recovery of polluted or burned soils.

## Introduction

Wildfires are increasing in severity and intensity (1), consuming nearly 4% of Earth’s land surface (∼464 M ha) every year (2), and altering the processes that govern the cycling and availability of carbon (C) and nitrogen (N) (3, 4). Wildfires also affect soil microbes (5, 6), altering their composition, function, and abundance. Fires reduce microbial richness and biomass, but they also promote specific taxa of pyrophilous “fire-loving” fungi (7), with uncertain effects on soil C and N cycling (4, 8). In particular, fires increase soil N availability (9) and generate complex charcoal-derived pyrogenic organic matter (PyOM) (10), which pyrophilous fungi can degrade. Despite these important biogeochemical roles, the genomic strategies and evolutionary trade-offs that enable pyrophilous fungi to survive and thrive post-fire are largely unknown.

Due to energetic costs and evolutionary constraints, trait trade-offs may limit fungal adaptation to post-fire biogeochemistry. Investment in certain traits, such as rapid growth or thermotolerance, could limit resource allocation to others like nutrient acquisition (11–13). Drawing from plant ecological theory and Grime’s C-S-R (Competitor-Stress tolerator-Ruderal) model (14), we previously proposed that pyrophilous microbes either excel at acquisition of post-fire resources such as high N or aromatic C in PyOM (competitors), thermotolerance (stress tolerators), or rapid growth in absence of competitors (ruderals) (12, 15). Investigating trait trade-offs provides insight into how fungi balance metabolic costs with ecological demands, which can result in a valuable framework for understanding and predicting microbial diversity and functions (11). Yet, empirical tests of these hypotheses remain scarce in microbial ecology. Trade-offs between stress tolerance, fast-growth, and C acquisition traits have been demonstrated in wood decaying fungi (16, 17), but not in other groups. Tests on cultures are a key step to determine whether these hypothesized trade-offs exist in pyrophilous fungi, with important implications for predicting global C and N cycling as fires continue to influence large tracts of the terrestrial biosphere.

Comparative genomics is an ideal approach to assess the genomic evolution and adaptations of pyrophilous fungi. Fungal genomes evolve through several mechanisms, including gene duplication, polyploidy, sexual reproduction, horizontal gene transfer (HGT), and transposable elements (TEs), all of which contribute to fungal functional and ecological diversity (18). Further, the unique prominent dikaryotic phase in Basidiomycota can lead to genomic novelty (19). Previous comparative analyses on several mushroom-forming taxa from Pezizales and Agaricales have revealed unique genomic signatures in pyrophilous fungi, notably, expansions in gene families related to stress response, carbohydrate metabolism, and secondary metabolite biosynthesis (20, 21). Yet, further research is needed to address potential trade-offs between these and other ecological or physiological traits.

We hypothesize that four key mechanisms–trait trade-offs, gene duplication, HGT, and sexual recombination–have driven the development of fire-associated traits in pyrophilous fungi. Trait trade-offs may have shaped fungal specialization, balancing investments in rapid growth, thermotolerance, and resource acquisition to maximize fitness in fire-disturbed ecosystems. Gene duplication combined with mutations, which is a major source of adaptation in fungi (22), likely plays a central role by expanding enzymatic gene families involved in PyOM degradation and N acquisition, enhancing metabolic versatility in post-fire environments. HGT, which can expand capacity and adaptation (23), may contribute to the acquisition of novel aromatic C degradation and N acquisition genes, enabling more efficient resource utilization and providing a competitive advantage in nutrient-limited post-fire soils. Finally, recombination during sexual reproduction could further increase genetic diversity by creating new combinations of alleles that optimize fungal responses to nutrient pulses and environmental stressors.

Here, we leveraged a large and diverse culture collection of pyrophilous fungi isolated primarily from post-fire mushrooms and soils across California (24) to identify genomic traits and trade-offs driving rapid post-fire colonization and persistence. We combined an *in-situ* growth assay with comparative genomics based on high quality, contiguous genomes of 18 pyrophilous fungi from orders of five Ascomycota and three Basidiomycota and found a dramatic trade-off between fast growth and genes encoding aromatic C degradation. Complementary transcriptomics analysis of four pyrophilous fungal isolates from four orders to PyOM amendment across time further suggested that energetic costs of transcriptional regulation contribute to observed trait trade-offs. Overall, we propose two major genome evolution strategies that underlie the pyrophilous fungal adaptation to post-wildfire environments: one relying on gene duplication and somatic mutation (e.g., in Eurotiales) and the other on sexual recombination (e.g., in Pezizales and Agaricales). Excitingly, we identified putative cross-kingdom horizontal gene transfer from bacteria to fungi as a rare third strategy (e.g., in Coniochaetales and Eurotiales) to enhance genetic diversity beyond the constraints of the gene pool. By generating new high-quality reference genomes for seven pyrophilous fungi, we advance understanding of fungal genome evolution, in general, as well as expand existing genomic research on pyrophilous fungi from two (20, 21) to eight fungal orders. Our findings support the hypothesis that trait trade-offs shape ecological function, with implications for global C and N cycling, and for creating microbial inoculants capable of breaking down polycyclic aromatic hydrocarbons that pollute many environments including burned soils.

## Results

### Genome assembly and gene annotation statistics

We generated 18 high-quality, contiguous fungal genomes from five Ascomycota orders (Coniochaetales, Eurotiales, Pezizales, Pleosporales, Sordariales) and three Basidiomycota orders (Agaricales, Geminibasidiales, Holtermanniales), which are all abundant in post-fire environments across a variety of fire-prone ecosystems (12, 15, 25), primarily from cultures isolated from burned soils and mushrooms following California wildfires (Table S1). We obtained 6.5-213 Gb of PacBio HiFi sequencing per genome with read N50 ranging from 5,429-10,028 bp providing 21-521X genome coverage (Table S2). Genome sizes ranged from 16.6 to 61.6 Mb, from 7 to 49 contigs, and contig N50 of 1.1 to 9.2 Mb (Table S3). Scaffolding with Masurca improved the contiguity of *Morchella eximia* (reduced number of contigs from 58 to 42), *Tricharina praecox* (from 52-41), and *Pholiota brunnescens* (from 53 to 49) assemblies. It also significantly improved contiguity of *N. discrete* E-DF3 (reduced number of contigs from 25 to 7) and *N. sp.* E-DF1 (from 41-23) genomes, where we used the chromosome-level assembly *of N. discrete* FGSC8579 as a reference.

We found that 16/18 of our genomes were monokaryotic, with 14 being Ascomycota, and the 2 Basidiomycota yeasts, *Basidioascus undulatus* and *Holtermanniella festucosa,* being functionally monokaryotic due to the low number of heterozygous SNPs (Table S4). However, the filamentous Basidiomycota, *Lyophyllum atratum* and *Pholiota brunnescens*, had a large number of heterozygous SNPs, suggesting that they are dikaryotic and must be treated as phased genomes (Supplementary Result 1 and Table S4). We also observed internuclear genome re-arrangement after the alignments of two haplotypes of *P. brunnescens* (Fig S1A) and *L. atratum* genomes (Fig S1B).

We also assembled 21 to 267 Kb-sized mitochondrial genomes with most isolates in a single contig (Table S5). Gene annotation guided by 2-15 million RNA-Seq paired-end reads per isolate enabled the identification 7,097-16,135 protein-coding genes per genome (Table S6). We sequenced the first genomes for 11 of our fungal isolates, with 6 providing completely novel reference genomes (Table S3, Figs. S2, S3), and provide more contiguous assemblies for 7 of them (20, 21, 26).

### Trade-off between aromatic C degradation gene composition and fast growth

We paired genomes with biophysical assays to test if enrichment of aromatic C degradation and N acquisition genes trade off with growth on Malt Yeast Extract Agar (MYA) and PyOM media made from wildfire charcoal. We observed large variations in the enrichment of aromatic C degradation and N acquisition genes among pyrophilous fungal orders where Eurotiales had the most genes, followed by Pleosporales, then Coniochaetales and the lowest in Sordariales, Pezizales and Agaricales (Results S2; Fig. S4). We found that in general the Sordariales and Pezizales had the fastest growth whereas Eurotiales had the slowest growth (Fig. 1A), with a significant positive correlation between hyphal extension on MYA and PyOM media (r=0.82, *P*=0.0001; Fig. 1B). There was a dramatic trade-off between fast growth and enrichment of post-fire resource acquisition genes as indicated by strong, significant negative correlations (r=0.53-0.78, *P*=0.0001-0.05) between MYA or PyOM HE rates and the number orthologues of most aromatic C degradation genes (Fig. 1B). The microbial pathways involved in the decomposition of aromatic C in PyOM primarily include naphthalene degradation, catechol meta- and ortho-cleavage, and protocatechuate meta- and ortho-cleavage (26–28). Fungi transform aromatic C into intermediates such as catechol, protocatechuate, and benzoyl-CoA, followed by conversion to secondary intermediates such as acetyl-CoA, succinyl-CoA, and pyruvate through the bKA pathway (29). Interestingly, the number of *pcaF* orthologues from the bKA pathway, *mhpD* from the catechol metacleavage pathway, and *catB* from the catechol orthocleavage pathway were negatively correlated solely with HE on PyOM media. Contrary to our hypothesis, the correlation between N acquisition orthologue number and HE on both media types was insignificant (Fig. 1B).

**Fig 1.**
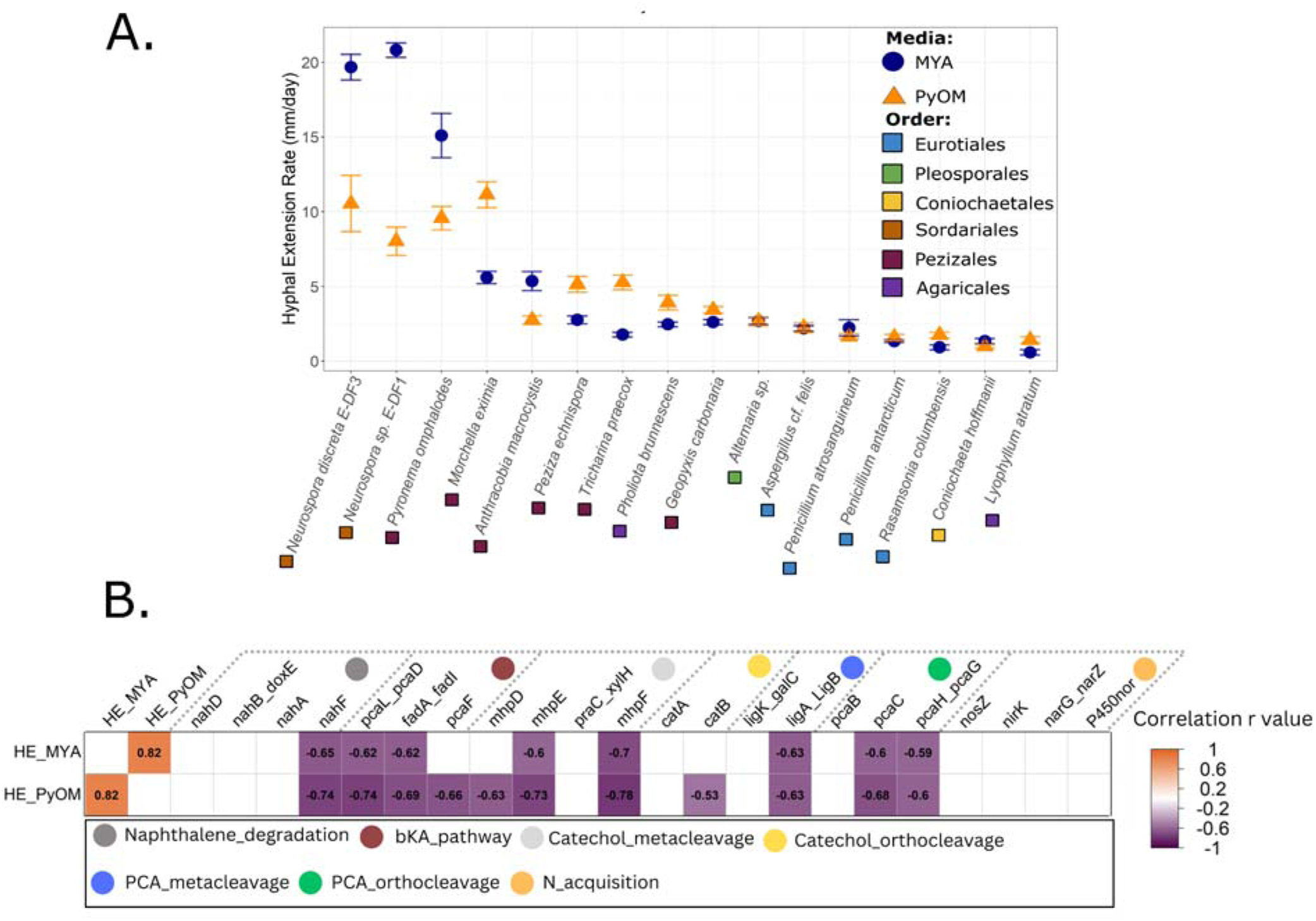
Hyphal extension of 16 pyrophilous fungi on Malt Yeast extract Agar (MYA) and Pyrogenic Organi Matter (PyOM) media **(A)** and pairwise Spearman’s rank correlation between the hyphal extension on MYA (HE_MYA) and PyOM (HE_PyOM) media and the number of aromatic C degradation and N acquisition orthologues in each isolate **(B)**. Error bars in panel A shows standard errors of mean. Number in the cells denote pair-wise correlation r values. Gene abbreviation in panel B; nahD: 2-hydroxychromene-2-carboxylate-isomerase, nahB_doxE: cis-1,2-dihydro-1,2-dihydroxy naphthalene, nahA: Naphthalene-1,2-dioxygenase subunit-alpha, nahF: salicylaldehyde_dehydrogenase, pcaL_pcaD: 3-oxoadipate-enol-lactonase, fadA_fadI:Acetyl-Co A acyltransferase, pcaF: beta-ketoadipyl-CoA-thiolase, mhpD: 2-keto-4-pentenoate hydratase, mhpE: hydroxy_oxovalerate_aldolase, praC_xylH: 4-oxalocrotonate_tautomerase, mhpF: acetaldehyde_dehydrogenase, catA: catechol-1,2-dioxygenase, catB: muconate cycloisomerase, ligK_galC: 4-hydroxy-4-methyl-2-oxoglutarate aldolase, ligA_LigB: Protocatechuate-dioxygenase, pcaB: 3-carboxy-cis,cis-muconate-cycloisomerase, pcaC: 4-carbox muconolactone-decarboxylase, pcaH_pcaG: Protocatechuate-3,4-dioxygenase, nosZ: nitrous-oxide-reductase, nirK: nitrite-reductase; narG_narZ: nitrate-reductase, P450nor: P450 nitric oxide reductase.

### Role of gene duplication in aromatic C degradation and N acquisition orthologues enrichment

We performed comparative genomic analysis using Orthofinder by adding 38 genomes of sister species with publicly available genomes to 16 pyrophilous isolates from our study within six fungal orders (Table S8). We excluded Geminibasidiales (*B. undulatus*) and Holtermanniales (*H. festucosa*) since they are the only genomes sequenced from those orders. We tested the contribution of gene duplication to gene enrichment by mining the Orthofinder output of the 54 genomes for the composition of 18 aromatic C degradation and 4 N acquisition orthologues (Table S9). We found significant (P < 0.05) phylogenetic signals for the enrichment of all aromatic C and N acquisition genes after normalization of gene count to the total proteome size of each genome (Table S7). Orthofinder partitioned duplication events to the current species (pyrophilous and their sister species) and their common ancestors (named arbitrarily by Orthofinder as N0-N51) in the species tree, providing insights into the duplication event history during evolution.

We inferred two distinct groups at the N2 common ancestor. The first group, containing the Eurotiales, Pleosporales, Sordariales, and Coniochaetales, had more aromatic C degradation orthologues, while the second group with Pezizales and Agaricales had fewer orthologues (Fig. 2). To test if more than one major duplication event led to the enrichment seen in the first group, we compared the common ancestors descended from N2. We found that the duplication rate at N6–the ancestor of Eurotiales, Pleosporales, Sordariales, and Coniochaetales–was 4X higher than at the concurrent node N5, which also descends from N2 and gave rise to the Pezizales and Agaricales. Meanwhile, N11 containing Sordariales and Coniochaetales, experienced a decline in duplication events, while N10 containing the Eurotiales and Pleosporales, and N18, the common ancestor of all contemporary Eurotiales maintained large duplication events. Within Eurotiales, the common ancestor of *Penicillium* at N36 had 10X higher duplication than the common ancestor of *Aspergillus* at N37. We found 36 genes-species combinations with significantly (P < 0.05) higher gene enrichment in pyrophilous than non-pyrophilous sister species (marked with blue rectangles in Fig. 2).

**Fig 2.**
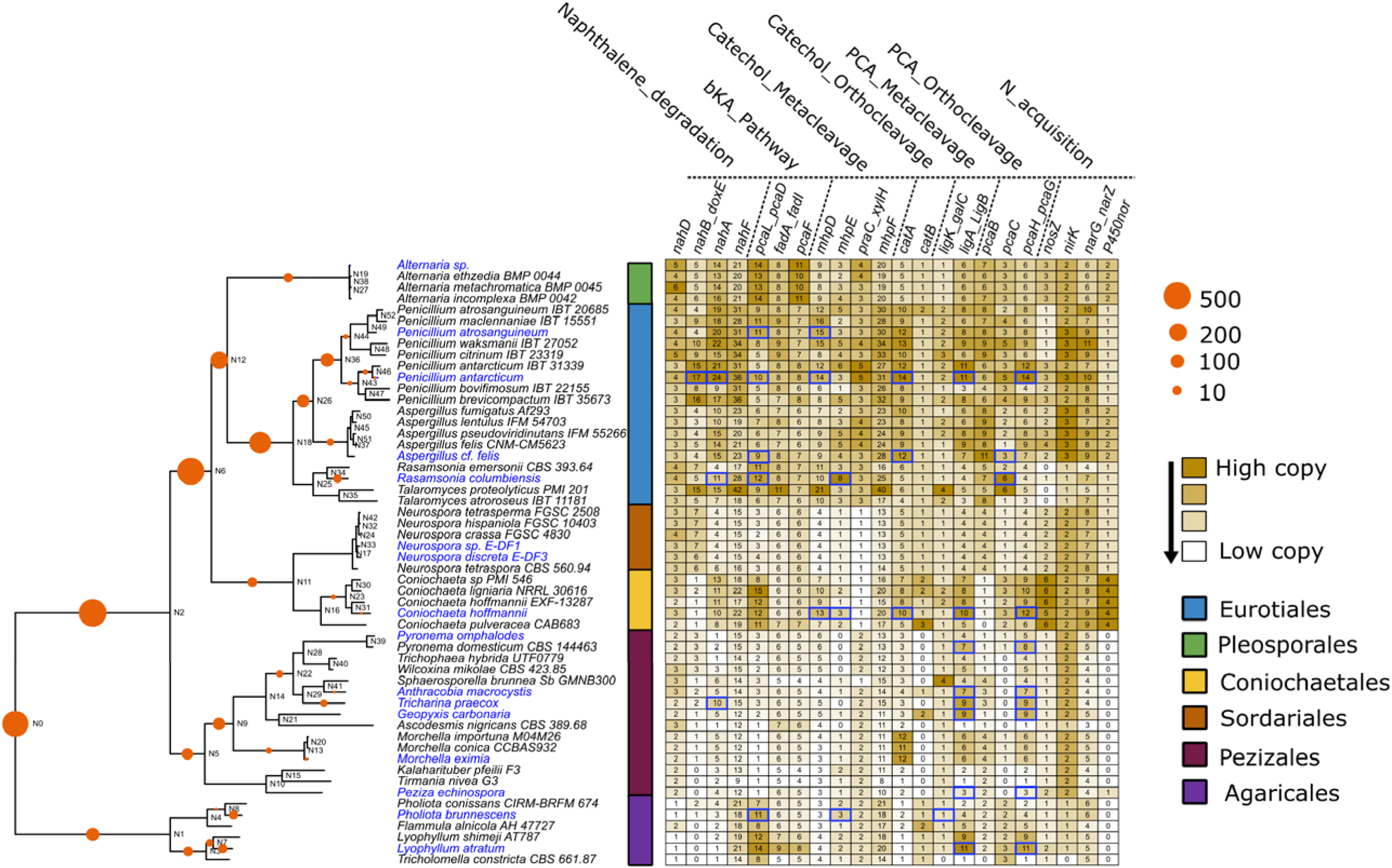
Number of duplication events (orange circles on the species tree) in the aromatic carbon (C) degradation (six pathways) and nitrogen (N) acquisition orthologues in 53 genomes used for constructing orthogroups and their common ancestors vs. the total number of orthologues of 18 aromatic C degradation and four N acquisition genes. The tree leaves printed in blue are 16 pyrophilous species sequenced in this study. The blue rectangles show significant (P<0.05) gene enrichment in pyrophilous vs. non-pyrophilous sister species inferred from Poisson distribution test. Gene name abbreviations: nahD: 2-hydroxychromene-2-carboxylate-isomerase, nahB_doxE: cis-1,2-dihydro-1,2-dihydroxy naphthalene, nahA: Naphthalene-1,2-dioxygenase subunit-alpha, nahF: salicylaldehyde_dehydrogenase, pcaL_pcaD: 3-oxoadipate-enol-lactonase, fadA_fadI: Acetyl-Co A acyltransferase, pcaF: beta-ketoadipyl-CoA-thiolase, mhpD: 2-keto-4-pentenoate hydratase, mhpE: hydroxy_oxovalerate_aldolase, praC_xylH: 4-oxalocrotonate_tautomerase, mhpF: acetaldehyde_dehydrogenase, catA: catechol-1,2-dioxygenase, catB: muconate cycloisomerase, ligK_galC: 4-hydroxy-4-methyl-2-oxoglutarate aldolase, ligA_LigB: Protocatechuate-dioxygenase, pcaB: 3-carboxy-cis,cis-muconate-cycloisomerase, pcaC: 4-carboxy muconolactone-decarboxylase, pcaH_pcaG: Protocatechuate-3,4-dioxygenase, nosZ: nitrous-oxide-reductase, nirK: nitrite-reductase; narG_narZ: nitrate-reductase, P450nor: P450 nitric oxide reductase.

For the two dikaryotic genomes, we found that 84% of aromatic C and N acquisition genes in *P. brunnescens* and 54% in *L. atratum* had copies in both haplotypes (Supplementary File 1), suggesting the contribution of both nuclei to the genomic enrichment. *L. atratum* had duplication in one of the haplotypes for 19% of aromatic C and N acquisition genes contributing further to the overall gene enrichment; however, no haplotypic duplication was observed for *P. brunnescens*. Internuclear gene presence/absence variation was 9% for *P. brunnescens* and 6% for *L. atratum*.

### Horizontal gene transfer involved in the higher enrichment of catA

Considering the phylogenomic relatedness of orders and reduced duplications events at N11 (Fig. 2), we expected both Coniochaetales and Sordariales to convergently evolve to lower number of aromatic C degradation and N acquisition orthologues than Eurotiales and Pleosporales. However, Coniochaetales species had notably more orthologues like catechol-1,2-dioxygenase (*catA*) from the bKA pathway than Sordariales, suggesting that gene duplication is not the only cause of aromatic C degradation and N acquisition orthologue enrichment.

We used six methods to test for HGT. First, we examined the gene trees of *catA* orthologues and identified OG0001245, an orthogroup carrying 63 *catA* orthologues. This lent support for HGT in the pyrophilous species *Coniochaeta hoffmannii*, *Penicillium atrosanguineum*, and *Rasamsonia columbiensis* (Fig. 3A), since each had two paralogues of *catA*, one of which clustered distantly from the orthologues of their sister taxa. Gene CH_06113.t1 from *C. hoffmannii* was clustered distantly from its paralogue CH_10993.t1, and close to an orthologue from *P. atrosanguineum* (PR_10622.t1). Gene PR_10622.t1 from *P. atrosanguineum* was clustered distantly from all the other Eurotiales orthologues including its paralogue PR_04517.t1. Gene RC_00189.t1 formed a distinct cluster with an orthologue from *R. emersonii* (XP_013325789.1) close to the Pezizales orthologues, while its paralogue RC_0097.t1 was clustered with the other Eurotiales orthologues. Due to the lack of identity by descent, we propose that CH_06113.t1, PR_10622.t1, and RC_00189.t1 were introduced through HGT.

**Fig 3.**
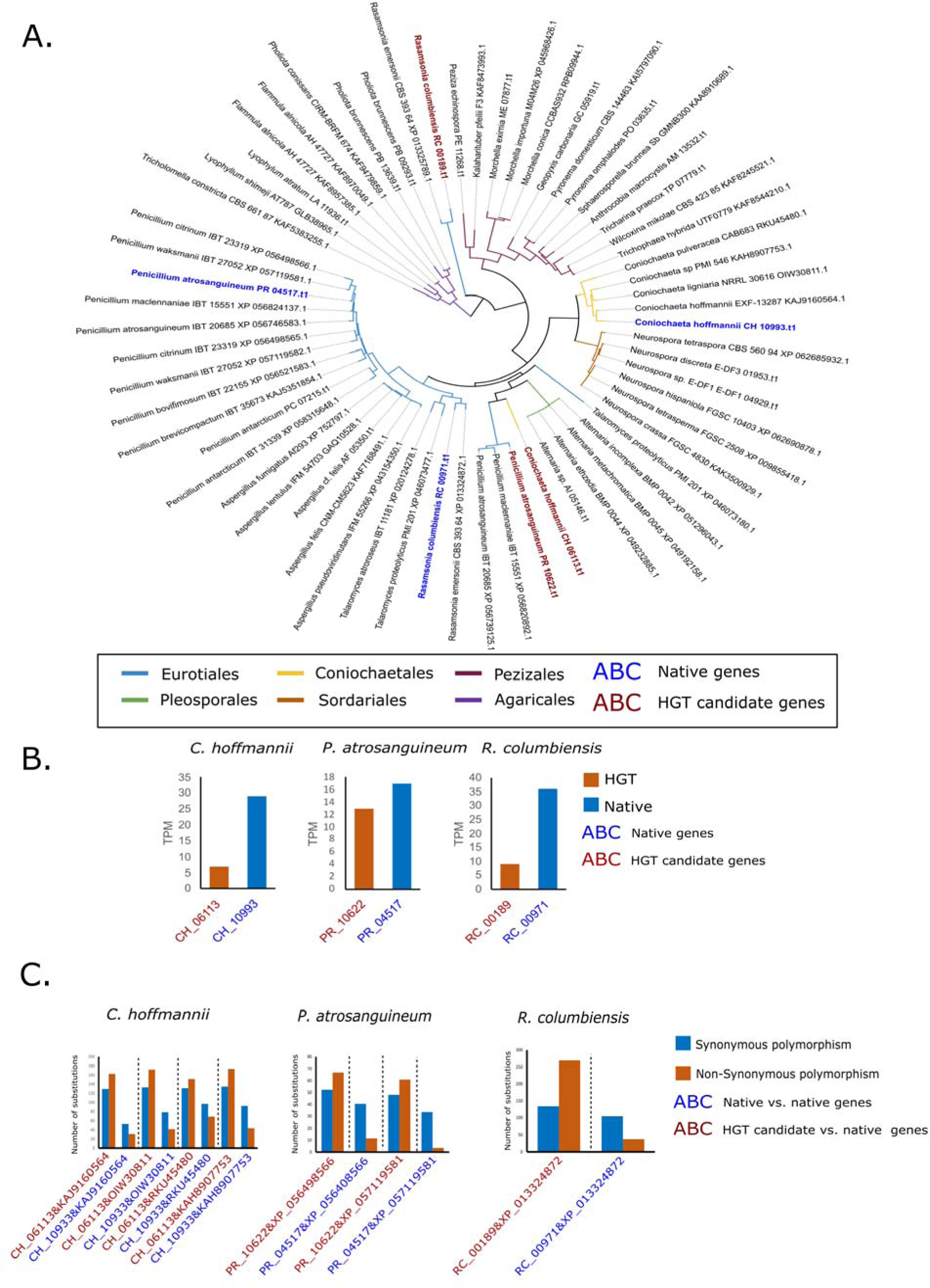
The Evidence supporting a horizontal gene transfer (HGT) event in OG0001245 orthogroup, described as catechol-1,2-dioxygenase (catA) in three pyrophilous species *Coniochaeta hoffmannii*, *Penicillium atrosanguineum*, and *Rasamsonia columbiensis*. Gene tree containing the 63 orthologues of OG0001245 orthogroups **(A)**, the transcription (transcript per million [TPM]) values of the candidate HGT genes and their native paralogues **(B)**, and the number of synonymous and non-synonymous nucleotide polymorphisms within the candidate HGT gene and their native paralogues vs. native orthologues of sister species **(C)**.

Second, we used multiple sequence alignment to align the protein sequences of the three HGT candidates and their native paralogues, and the resulting phylogenetic tree supported the distinction between the HGT candidates and their native paralogues (Fig. S5). We found a large deletion in CH_06113.t1, making it distinct from all the other five proteins (i.e., its native paralogue CH_10933.t1). Third, since previous research indicated that HGT introduced genes had lower transcription than their native counterparts (30), we examined their transcription levels and found that all three candidate HGT genes had lower transcripts per million (TPM) values than their native counterparts (Fig. 3B). Fourth, we noted that previous research found that HGT genes accumulate higher nucleotide polymorphisms than their native paralogues (31). We then use variation in sister taxa orthologues as a baseline for comparison, and we found higher numbers of nucleotide polymorphisms between orthologues of sister taxa and HGT candidates than their native paralogues (Fig. 3C). The difference was often larger for non-synonymous nucleotide polymorphisms (polymorphisms leading to protein amino acid changes).

We then search for the co-localization of Transposable Elements (TE) with HGT candidates and found a Ty3-line (Gypsy) Long Terminal Repeat (LTR) retrotransposon of ∼18 kb spanning CH_06113.t1 and the two neighboring genes (Fig. S6). Finally, for our sixth method, we used a complementary analysis of pyrophilous bacterial genomes to examine the possibility of cross-kingdom HGT from bacteria to fungi. The BLAST search against the Swiss-Prot database described CH_06113.t1 as “similar to hqdA Intradiol ring-cleavage dioxygenase hqdA (*Aspergillus niger*)”, and one of the neighboring genes as “similar to tftE Maleylacetate reductase (*Burkholderia cepacia*)”. When the protein sequence of CH_06113.t1 was used as an inquiry for NCBI BLASTP search against prokaryotes gene bank, we found several orthologues. We retrieved the protein sequences of the top 20 blast hits based on the percentages of sequence identity and query coverage and combined with the protein sequences of 20 fungal orthologues, and an orthologue we found on a conjugative plasmid of *Noviherbaspirillum soli,* a pyrophilous bacterial species we isolated from the same location as *C. hoffmannii*, for phylogenetic analysis. We found that the bacterial orthologues formed a distinct cluster from those of fungi in the phylogenetic tree, except for the *N. soli* plasmid orthologue that clustered with CH_06113.t1, suggesting a candidate HGT from *N. soli* to *C. hoffmannii* (Fig. S7). Despite closest phylogenetic relatedness, we still found phylogenetic distances between the plasmid and the candidate HGT genes (Fig S7), which could suggest an ancient transfer and an independent accumulation of mutations in fungi and bacteria over the course of evolution.

### Role of sexual reproduction in high variation in aromatic C and N acquisition genes

We tested if sexual recombination contributed to the genetic variation in aromatic C and N acquisition genes in pyrophilous fungi. To achieve that we measured nucleotide substitution rates across taxa, under the expectation that lineages with regular sexual reproduction should exhibit higher substitution rates than predominantly asexual species. We examined synonymous (S) and nonsynonymous (NS) nucleotide substitution rates separately to distinguish neutral mutation rates from adaptive selection pressures to determine whether variation in the nutrient acquisition pathways arose from selection for metabolic specialization (e.g., in aromatic C degradation) or neutral diversification processes. To facilitate the visualization of results, we selected one orthogroup from each of the examined pathways (Table S9).

We found significant (P < 0.05) phylogenetic signal for the number of nucleotide substitutions normalized to the total proteome size of all the examined orthogroups (Table S10). Eurotiales had consistently high numbers of S and NS substitutions across all pathways (Fig. 4); however, Eurotiales had the lowest overall difference between pyrophilous and non-pyrophilous sister species, in line with strong phylogenetic signal. Pezizales species exhibited comparable and sometimes higher S and NS substitutions than Eurotiales, despite having remarkably lower gene duplication events (Figs. 2; S4). Two Pezizales isolates, *Peziza echinospora* and *Geopyxis carbonaria*, ranked highest in the substitution number of the catechol metacleavage and naphthalene degradation orthogroups (Fig. 4).

**Fig 4.**
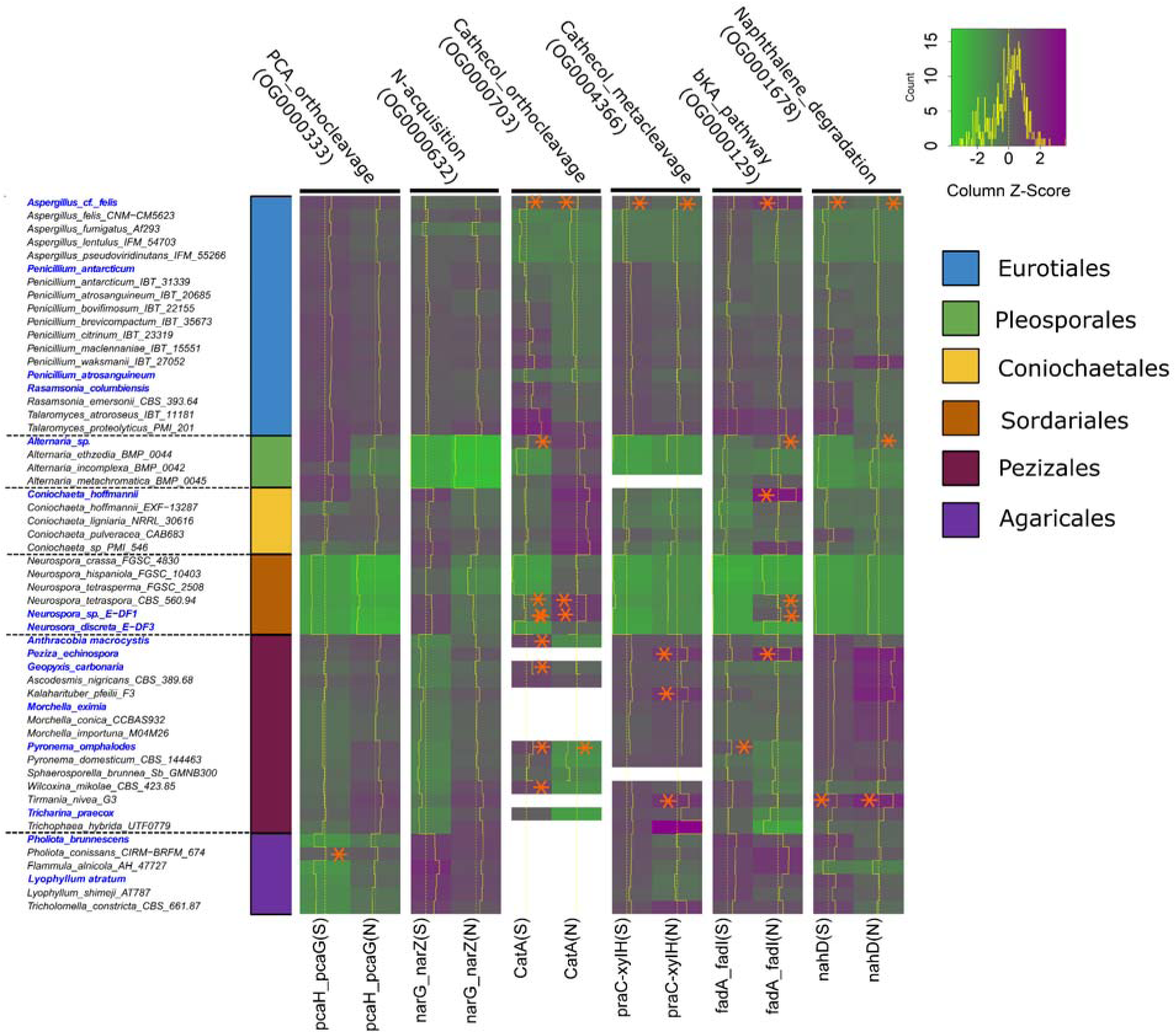
The number of synonymous and nonsynonymous nucleotide substitution in six orthogroups representing five aromatic C degradation and the N acquisition pathways in 53 genomes used for orthogroup construction. Taxa in blue text were sequenced in this study. The dash line in each heatmap cell shows the median number of nucleotide polymorphisms across all taxa for each orthogroups and the solid line is that of each taxon. The heat map color scale represents nucleotide substitution counts for each orthologue, with green indicating lower and purple indicating higher frequencies. The labels at the top of the heat map denote the names of selected orthogroups from each pathway, while those at the bottom indicate gene names and whether the substitutions are synonymous (S) or nonsynonymous (N). The stars denote significantly higher (P < 0.05) number of genetic variants in pyrophilous than their non-pyrophilous sister species inferred from Poisson distribution test. Orthogroups selected were described as Protocatechuate-3,4-dioxygenase (*pcaH_pcaG*), nitrate reductase (*narG-narZ*), catechol-1,2-dioxygenase (*catA*), 4-oxalocrotonate tautomerase (*praC-xylH*), Acetyl-Co A acyltransferase (*fadA_fadI*), and 2-hydroxychromene-2-carboxylate isomerase (*nahD*). White cells in heat-map indicated absence of genes in the corresponding taxa, impeding the calculation of the substation rate.

To distinguish pyrophilous adaptations from phylogenetic relatedness, we compared pyrophilous fungi and their non-pyrophilous sister taxa and found that some pyrophilous fungi exhibited significantly (P < 0.05) higher nucleotide substitutions in aromatic C degradation genes than their sister taxa. We found that across 38 orthogroups-species combinations (shown with asterisk in Fig. 4) where nucleotide polymorphism differed between pyrophilous and non-pyrophilous species, with 29 showing higher nucleotide variation in pyrophilous species. *Aspergillus cf. felis* had significantly higher S and NS substitutions in the catechol orthocleavage, catechol metacleavage, and naphthalene degradation orthogroups than all the other sister *Aspergillus* species, and *Alternaria* sp. had higher substitutions in the catechol orthocleavage, bkA pathway, and naphthalene degradation orthogroup than all the other *Alternaria* species (Fig. 4). *Coniochaeta hoffmannii* had the highest NS substitutions in the bKA pathway orthogroups within the Coniochaetales and also ranked highest among all 53 taxa.

### Dikaryotic internuclear variation contributes to nitrate-reductase (narG-narZ) variation

The dominant dikaryotic phase common in Basidiomycota where two genomes exist in two separate nuclei in each cell allows them to take advantage of diploid genomes (e.g., increased genetic diversity and protection against deleterious mutations) while reducing the tension between unrelated haploid genomes (32). We leveraged our high-quality assemblies to establish phased genomes for the dikaryon *L. atratum* to test the contribution of internuclear variation to higher nucleotide substitution observed in nitrate-reductase (*narG-narZ*) genes (Fig. 4). We found a high number of substitutions in the Agaricales OG0000632 orthogroup described as *narG-narZ*. We selected an OG0000632 orthologue of *Lyophyllum atratum* (LA_01440.t1) and aligned two haplotypes representing the sequences of two nuclei and found 16 nucleotide polymorphism sites in the exonic regions (Fig. S8). We then aligned RNA-seq reads to one haplotype as a reference and observed reads derived from both haplotypes at a nearly 1:1 ratio at all 16 polymorphism sites, indicating that the two haplotypes were transcribed in equilibrium. This suggests that internuclear variation drove the higher number of substitutions in *narG-narZ* in *L. atratum*.

### Transcriptome response of selected pyrophilous fungi to PyOM treatment

We performed transcriptomics at 0, 2, 4, 24 and 48 hours after PyOM amendment to assess how PyOM affected gene transcription across time for 4 isolates. We selected *Coniochaeta hoffmannii*, *Penicillium atrosanguineum*, *Morchella eximia,* and and *Pholiota brunnescens* representing four orders with distinct levels of aromatic C degradation and N acquisition gene enrichment. We found that orthologues of 14 genes in *C. hoffmannii* (Fig. S9), 21 in *P. atrosanguineum* (Fig. S10), 17 in *M. eximia* (Fig. S11), and 15 in *P. brunnescens* (Fig. S12) were differentially transcribed in response to PyOM treatment. There was a common set of aromatic C degrading genes showing differential transcription in all fungi, for example the catA orthologues. In contrast, transcription of certain gene orthologues like *nahB_doxE* only positively responded to PyOM only in *P. atrosanguineum* (Fig. S10) and *M. eximia* (Fig. S11), despite being present in all four species (Fig. 2). In terms of N acquisition, *narG-narZ* was induced in all four species and *nirK* was induced in all but *C. hoffmannii*.

## Discussion

We provided an experimental test of ecological theory to reveal a fundamental trait trade-off between ability to grow fast and acquire post-fire resources in pyrophilous fungi. This genomic balancing act underpins the selective advantage of carrying larger suites of aromatic C degradation genes in wildfire-impacted environments. Our findings suggest that gene duplication and mutations represent one route for generating variation within aromatic C degradation and N acquisition genes in pyrophilous fungi (e.g., in Eurotiales), while recombination via sexual reproduction and internuclear variation offer an alternative mechanism (e.g., in Pezizales and Agaricales). We also show evidence that ancient bacterial to fungal cross-kingdom HGT contributes to genetic novelty in PyOM degradation in pyrophilous fungi (e.g., in Coniochaetales and Eurotiales). Finally, we found stringent transcriptional regulation of aromatic C and N acquisition genes in four distantly related pyrophilous fungi, showing low baseline transcription in regular media but strong induction following PyOM treatment. This finding suggests that the energetic costs of transcription contribute to observed trait trade-offs and that transcriptional responses to PyOM are shaped less by phylogeny and more by shared ecological adaptation to post-fire environments. We integrate comparative genomics, transcriptomics, and growth assays to link specific genetic changes with ecological outcomes in a diverse array of understudied pyrophilous fungi that may have exceptional potential as fungal inoculants for ecosystem recovery after disturbances.

### Microbial trait trade-offs at genomic levels

Combining hyphal extension assays with high-quality genomes of 16 filamentous pyrophilous fungal isolates, we found large, significant trade-offs between fast growth and enrichment for aromatic C degradation genes. For instance, the Ascomycota order Pezizales grew fast at the expense of fewer aromatic C breakdown orthologues, while Eurotiales grew slowest but was enriched for aromatic C degradation. Surprisingly, we found no trade-offs between fast growth and enrichment of genes for N acquisition, which may reflect that inorganic N is so prevalent across post-fire ecosystems (cite) that it is not necessary to invest heavily in acquiring it, or alternatively that efficient N scavenging is a widespread evolutionary pressure across most ecosystems (33). In contrast, the recalcitrance of PyOM requires highly specialized enzymatic machinery (34), creating a stronger metabolic cost driving the trade-off with growth. The fast-growth-aromatic C acquisition trade-off observed here, along with genetic and copy number variation within the aromatic C degradation and N acquisition orthologues, supports the ideas of functional diversity and specialization among pyrophilous fungal lineages. The rapid growth of Pezizales species corresponds to their early successional role in post-fire environments, coinciding with a surge of bioavailable nutrients from necromass (15, 35). Similarly, the comparatively slower growth of Eurotiales aligns with their late successional role, where enrichment of aromatic C-degrading genes likely facilitates resource acquisition from difficult to degrade PyOM (36) Many Eurotiales are known for their rapid colonization of substrates (37), which made their slower growth unexpected; however, it might be specific to the pyrophilous Eurotiales species we examined, as evidenced by higher enrichment of aromatic C genes in pyrophilous Eurotiales and their sister species than other Eurotiales. This variation in growth rates among Eurotiales highlights the diverse ecological strategies within the order, ranging from fast-growing opportunists to more specialized, slower-growing species adapted to specific niches like post-fire environments.

### New pyrophilous taxa as candidate contributors to aromatic C degradation and N acquisition

Our results suggest that three Ascomycota orders–Eurotiales, Pleosporales, and Coniochaetales–are likely major players in aromatic C degradation due to gene enrichment. We know of only two prior studies examining pyrophilous fungal genomes that focused on six isolates of Pezizales and three of Agaricales that all formed mushrooms on burnt soil (20, 21). In contrast, Eurotiales, Pleosporales, and Coniochaetales do not form macroscopic fruiting bodies but are commonly identified in microbiome studies of burned soils (12, 15, 25) and we isolated them from burned soil slurries (24). The Eurotiales isolates we studied included *Rasamsonia columbiensis,* which is a thermotolerant species (37) that has been isolated from house dust (38), *Penicillium atrosanguineum,* which has been isolated from forest soil (39), *P. antarcticum*, which was previously isolated as an endophyte from Antarctica native plants (40), and *Aspergillus fellis,* also previously isolated as an endophyte (41) and from rocks (42). Endophytic fungi may be potent decomposers of aromatic C due to their ability to decompose the host’s dead cell wall and penetrate host plant barriers while establishing endophytic relationships (43). The endophytic lifestyle could be linked with post-fire survival by providing an escape mechanism, as internal colonization of plant tissues, especially roots, offer insulation from extreme heat and flames, allowing fungi to persist and subsequently colonize burned soil (44, 45). Pleosporales, such as *Alternaria*, are adapted to a wide range of ecological niches and are ubiquitous saprobes, wide-host-range plant pathogens, and plant endophytes (46). *Coniochaeta hoffmannii* is also known as a wood-decaying soil fungus and a plant pathogen that was previously isolated from a car fuel reservoir and can degrade polypropylene plastic (47). Eurotiales, Coniochaetales, and Pleosporales pyrophilous species, characterized by enriched aromatic C degradation and N acquisition genes and extremophilic adaptations to environments like Antarctic soils, fuel reservoirs, and pyrolyzed substrates, exhibit dual capacities critical to post-wildfire recovery–degradation of fire-derived aromatics (e.g., polypropylene-like pyrolyzed compounds), and tolerance of post-fire abiotic stressors (e.g., desiccation and UV exposure). Their genomic toolkit, refined through adaptation to post-fire conditions, positions these fungi as key mediators of C and N cycling in burned soils, with potential applications in restoring fire-disturbed ecosystems. In contrast, the Basidiomycete pyrophilous fungi did not have as many genes for aromatic C degradation, which may reflect their specialization in lignin degradation (48).

We assembled contiguous genomes for two basidiomycete yeasts, *Holtermanniella festucosa* from Holtermanniales and *Basidioascus undulatus* from Geminibasidiales, which were both genomic orphan orders. These species are novel pyrophilous fungi and have been repeatedly identified in previous studies as dominant soil colonizers following wildfire (12, 15, 25, 49). However, due to the absence of closely related reference genomes, we were unable to include these isolates in our comparative genomic analyses.

### Gene duplication, an adaptation strategy to post-fire environments in pyrophilous fungi

Our study revealed that ancient gene duplication plays a crucial role in shaping the current composition of aromatic C degradation and N acquisition genes in pyrophilous fungi, likely as an adaptation to frequent exposures to post-fire environments. Gene duplication is a well-established mechanism for creating genetic variation and functional innovation across different forms of life and especially within Kingdom Fungi, where five large gene duplication bursts have been identified (50). Among our isolates, gene duplications decreased throughout evolutionary history, but this decline was slower for the common ancestors of the current pyrophilous species in the Eurotiales, Pleosporales, Sordariales, and Coniochaetales orders. Eurotiales species showed the slowest decline, maintaining the largest repertoire of aromatic C degradation genes in their contemporary forms. This slow decline leading to high load of aromatic C degradation genes highlights the significant role that Eurotiales likely play in aromatic C degradation, at least in fire disturbed chaparral soil, where our pyrophilous Eurotiales species were isolated (24). A previous study found a correlation between gene copy numbers of ring-hydroxylating dioxygenase in soil samples and degradation efficiency, highlighting the quantitative nature of microbial aromatic C degradation (51). Gene duplication thus provides novel genes for higher expression and raw materials for genomic novelty for functional diversification. While gene duplication affecting C and N metabolism is a common evolutionary strategy in other fungi (22), our findings underscore its particular importance for pyrophilous fungi supported by the significant enrichment of some genes in pyrophilous vs. non-pyrophilous sister species. The higher enrichment in pyrophilous fungi despite strong phylogenetic signal likely represents a specific adaptation to unique post-fire C and N resources. However, we observed differing duplication frequencies between aromatic C and N acquisition genes across pyrophilous fungal species supported with strong phylogenetic signal. This suggests lineage-specific variations in employing gene duplication as an adaptive strategy post-fire. Previous studies also suggested that pyrophilous fungi are phylogenetically conserved, with the Eurotiomycetes and Pezizomycetes having more pyrophilous fungi than expected by chance (12).

### Enhancing genetic variation in aromatic C and N acquisition genes through ancient HGT

Here, we provide for the first time compelling evidence for an ancient HGT of a *catA* gene from bacteria to *C.* 3/18 pyropilous fungi, *hoffmannii, R. columbiensis* and *P. atrosanguineum.* HGT is rare in fungi, making its occurrence in multiple pyrophilous species surprising and suggesting that the extreme, stress-intensive post-fire environment may promote such events, analogous to how the gut environment facilitates HGT (52). Our phylogenetic analyses using the *catA* orthologues in fungi, bacteria, and a plasmid gene we discovered through sequencing the genome of the pyrophilous bacterium *N. soli*, suggested distinction between most bacterial and fungal sequences. It is likely due to an ancient HGT event subjected to distinct evolution trajectories in fungi or the conjugation of a *N. soli* plasmid gene that has a protein sequence distinct from all *catA* in the bacterial kingdom. The former is supported by the presence of *catA* HGT events in distantly related orders of Coniochaetales and Eurotiales, and the latter by the clustering of plasmid-born *catA* with the fungal HGT candidate gene. It is possible that this gene was transferred into their common ancestor and subsequently lost in some descended taxa possibly due to fitness penalties. HGT from bacteria to fungi is rare but is increasingly recognized as a significant mechanism for fungal genome evolution and adaptation (53). Notable examples include the transfer of bacterial glycosyl hydrolases to Neocallimastigomycota rumen fungi, enhancing their ability to break down plant material (54). The HGT events in aromatic C degradation genes like *catA* may have conferred a novel function allowing pyrophilous fungi to outperform native soil communities in acquiring C from PyOM, which might be tested with future research generating a knockout of the HGT *catA* gene in pyrophilous fungi like *C. hoffmannii*. The cross-kingdom HGT of aromatic C acquisition genes is a novel mechanism of adaptation in pyrophilous fungi to post-fire environments, where higher genetic variation in aromatic C degrading genes is likely advantageous.

### Contribution of sexual recombination to pyrophilous fungal genetic variation

Sexual recombination generates novel allelic combinations and increases nucleotide diversity, whereas strictly asexual species accumulate variation more slowly, potentially limiting their genetic diversity despite multiple gene duplications. The number of nucleotide polymorphisms varied substantially across pyrophilous orders and surprisingly did not linearly increase with the number of gene duplications. Pezizales exhibited comparable, and in some cases, higher nucleotide polymorphisms than Eurotiales, despite having significantly fewer gene copies. Differences in sexual reproduction frequency, which is rare among Eurotiales species, with some reproducing strictly asexually (37), but common among pyrophilous Pezizales species (20), may suggest two distinct mechanisms for maintaining diversity in pyrophilous fungi. In fungi with limited sexual reproduction, like Eurotiales, genetic variation could be maintained through gene duplication followed by somatic mutations. In contrast, genetic variation might balance through sexual recombination in lineages like Pezizales with frequent sex. The high genetic variation observed in aromatic C degradation and N acquisition genes could allow pyrophilous Pezizales species such as *Pyronema omphalodes* to access various bioavailable C and N resources released from necromass immediately post fire, explaining the rapid colonization of burned soil by *Pyronema* (15, 35). The slower-growing Eurotiales pyrophilous species appear to take over later in succession (15), likely due to their higher gene copies which enable them to acquire more complex aromatic C and N resources that are inaccessible to pyrophilous Pezizales.

Despite strong phylogenetic signal, we observed significant differences between pyrophilous and non-pyrophilous sister species in the number of nucleotide substitutions of some genes, particularly for *Aspergillus cf. felis*, suggesting that stressful post-fire habitats can promote elevated genetic variation as a strategy for adaptation (55). We identified that inter-nuclear nucleotide variation contributed to the total composition of aromatic C and N acquisition genes for two dikaryotic basidiomycetes *P. brunnescens* and *L. atratum*. We also identified internuclear variation in *L. atratum* associated with a nitrate-reductase (*narG-narZ*) gene that exhibited unexpectedly high genetic variation. This finding further supports the role of sexual reproduction and dikaryon formation in Basidiomycota as a mechanism for enhancing genetic variation in resource acquisition genes among pyrophilous fungi. Together, these results underscore multiple evolutionary strategies employed by pyrophilous fungi to maintain genetic diversity and adapt to the dynamic nutrient landscapes of post-fire environments.

### Transcriptional changes in aromatic C and N acquisition genes in response to PyOM treatment

In addition to the RNA-sequencing for genome annotation, we performed differential gene transcription across time after PyOM amendment. We selected 4 isolates, *C. hoffmannii, P. atrosanguineum*, *M. eximia*, *P. brunnescens*, representing four different orders ranging from lower to higher gene enrichment for aromatic hydrocarbons and N acquisition. We observed strong transcriptional activation of key aromatic C degrading genes after PyOM treatment, indicating that expressing these pathways is metabolically costly and supporting the growth trade-off demonstrated in our bioassays Our results align with previously published results suggesting the charcoal modulating genes were differentially transcribed in two pyrophilous sister species *Pyronema omphalodes* and *P. demesticum* after PyOM treatment (20). We found aromatic C and N acquisition genes commonly transcribed in four species, suggesting partial conservation of response to PyOM in pyrophilous fungi. This pattern indicates that the transcriptional response is driven less by phylogenetic relatedness and more by shared ecological adaptation to post-fire environments. Similar patterns have been reported in soil fungi, where enzyme activity and nutrient acquisition functions remain conserved across communities despite high taxonomic variability (56).

### Conclusions

Our study advances fungal genomics by expanding the genomic resources for pyrophilous fungi, including several previously uncharacterized lineages. Leveraging our large and diverse culture collection of pyrophilous fungi (cite), we use high-quality genomes and complementary transcriptomics and bioassays to conduct one of the first experimental tests of trait trade-offs in fungi. We reveal that pyrophilous fungi exhibit a striking trade-off between rapid growth and the enrichment of aromatic C degradation genes, an adaptation that likely confers a selective advantage in fire-affected soils dominated by PyOM. We also found that this trade-off did not exist for N genes, suggesting that there is less metabolic cost to invest if N acquisition machinery. Our study suggests that at least two major genome evolution strategies drive the adaptation of pyrophilous fungi to post-wildfire environments: one relying on gene duplication and somatic mutation (e.g., in Eurotiales) and the other on sexual recombination (e.g., in Pezizales and Agaricales). HGT events, as examined in *C. hoffmannii*, *R. columbiensis* and *P. atrosanguineum*, is likely a third rarer strategy enhancing genetic diversity in pyrophilous fungi beyond the gene pool barrier. Future research should investigate how these distinct evolutionary strategies influence interactions among pyrophilous fungi, prioritizing the study of newly identified species with enriched aromatic C and N acquisition genes. Collectively, our findings expand the frontier of pyrophilous fungal genomics, advance our ability to predict their roles in global C and N cycling, and identify functionally robust pyrophilous isolates that could serve as microbial inocula for post-fire soil restoration.

## Materials and Methods

### Fungal isolates

We leveraged our collection of pyrophilous fungi isolated primarily from post-fire mushrooms and soils across California. We included 18 pyrophilous fungal isolates from across two phyla and 8 orders (Table S1). We selected these isolates because they were abundant across a variety of fire-prone ecosystems (7, 12, 15, 25) and we expected them to exhibit a diversity of pyrophilous trait strategies (12).

### High Molecular Weight (HMW) DNA Extraction and PacBio HiFi Sequencing

We extracted HMW DNA from fungal tissue with either Qiagen Genomic Tip 100/G (Qiagen, Hilden, Germany) or using a CTAB:Phenol:Chloroform method (57) if extraction with Qiagen Tips led to low DNA concentration (Table S1). See Methods S1 for fungal tissue preparation and DNA extraction details.

We then submitted DNA to the University of California Irvine Genomics Research and Technology Hub (UCI GRTH) for HiFi SMRT Bell Library Preparation and PacBio HiFi Sequencing. The sequencing library was prepared using SMRTbell template prep kit version 3.0 with the overhang barcoded adapters (PacBio, Menlo Park, USA) and sequenced on the Sequel IIe™ System (PacBio, Menlo Park, USA), except for *Neurospora sp.* E-DF1, *Neurospora discreta* E-DF3, and *Tricharina praecox*, which were sequenced using the Revio™ system (PacBio, Menlo Park, USA). Samples were multiplexed to achieve the genome coverage ≥ 20X.

### Genome Assembly Pipeline

We generated draft genome assemblies for each isolate using Hifiasm v.0.16.1-r375 (58), IPA v.1.8.0 with --no-phase command for haplotype genomes (https://github.com/PacificBiosciences/pbipa), hiCanu v.2.2 (59), Peregrine v.0.4.13 with reduction factor of 4 specified with -r 4 and window size of 64 specified with -w 64 (https://github.com/cschin/peregrine-2021?tab=readme-ov-file), and Flye v.2.9-b1774 (60). For phased assemblies of hiCanu and Hifiasm, the represented primary haplotype for haploid dikaryotic genomes were obtained using purge-dups v.1.2.6 with default parameters (https://github.com/dfguan/purge_dups). We obtained the assembly metrics using QUAST software v.5.2.0 (61), and used BUSCO software v.5.4.3 to assess assembly completeness (62). We used the most contiguous and complete assembly as a reference for scaffolding with the Masurca software v.4.1.0 SAMBA tool (63) with raw HiFi reads and then with the other inferior draft assemblies or available chromosome-level genome assemblies as inputs. We used the chromosome-level assembly of *N. discreta* FGSC8579 as reference (https://genome.jgi.doe.gov/portal/pages/projectStatus.jsf?db=Neudi1&utm_source=chatgpt.com) for scaffolding *N. discreta* E-DF3 and *N.* sp. E-DF1 genomes(63). We visualized the alignment of the HiFi reads to genome at the genomic loci where two contigs were merged through scaffolding in Integrative Genome Viewer (64). Breaks were introduced if read support was insufficient. Finally, we identified and corrected the miss-assemblies using CRAQ software v.1.0.9 (https://github.com/JiaoLaboratory/CRAQ). We found telomere repeat sequences with the FindTelomeres script (https://github.com/JanaSperschneider/FindTelomeres). We assembled the mitochondrial genome using get_organel software v.1.7.7.1 (https://github.com/Kinggerm/GetOrganelle) with the Hifiasm assembly graph as an input We identified heterozygous single nucleotide polymorphisms (SNPs) between the haplotypes of the dikaryotic genomes and phased them using HapCUT2(65). We used the number of heterozygous SNPs as a proxy for the extent of internuclear genetic variation in the basidiomycete genomes assumed to have haploid dikaryotic genomes. We also identified internuclear genomic rearrangement in haploid dikaryotic genomes by genome-wide alignment between the contigs obtained for two haplotypes using MUMmer v.4.0.1 (66).

### RNA extraction and sequencing

We grew the 18 fungal isolates in three conditions: in malt yeast broth 1) at ambient temperature 2) in ambient temperature followed by a 1 hr 40 °C heat-shock, or 3) on PyOM agar plates covered with cellophane sheets at ambient temperature. In addition, we selected a subset of 4 fungal isolates (*C. hoffmannii*, *P. atrosanguineum*, *M. eximia*, and *P. brunnescens*) to test the transcriptional changes in aromatic C and N acquisition genes after PyOM amendment. The four fungal isolates were first grown in MYB medium (Methods S1) for 5 days, and the resulting biomass was harvested and exposed to PyOM liquid media (24). Samples from three biological replicates per condition were collected at 0 (before PyOM amendment), 2, 4, 24, and 48 h post PyOM treatment for RNA extraction and mRNA sequencing. Differential transcription analysis was performed using DESeq2 (67), with all time points compared against the 0 h for the calculation of fold change in transcription. Statistical significance of differential transcription was determined in DESeq2 using adjusted p-values (Benjamini–Hochberg false discovery rate < 0.05). We extracted total RNA with the E.Z.N.A.® Fungal RNA Mini Kit (Omega Bio-Tek, Norcross, U.S.A.) following manufacturer’s protocol. See Methods S2 for method used for the RNA quality control. We prepared mRNA Illumina libraries using the Illumina TruSeq® RNA Sample Preparation v. 2 Kit (Illumina, San Diego, USA) and sequenced on the NovaSeq 6000 system (Illumina, San Diego, USA) at the UCI GTRH for annotation and at the U.S. Department of Energy Joint Genome Institute for the PyOM treatment experiment.

### Gene and TE Annotation

We used RNA-seq data to annotate the protein-coding genes with the FunGAP pipeline (68) after trimming sequencing adapters with TrimGalore v.0.6.10 (https://github.com/FelixKrueger/TrimGalore), and filtering to retain only reads with a Phred quality score > 30 and length of ≥ 50 bp. We assigned putative sequence descriptions to protein sequences predicted by FunGAP by searching with NCBI-BLASTP v.2.16.0 (https://blast.ncbi.nlm.nih.gov) against the reviewed UniProtKB database (Swiss-Prot, downloaded in Aug 2023). For each protein, we extracted the InterPro protein families and Gene Ontology (GO) terms (69) using InterProScan v. 5.62-94.0 (https://interproscan-docs.readthedocs.io/en/latest/) and the Kyoto Encyclopedia of Genes and Genomes (KEGG) ontology terms using kofam_scan v.1.3.0 (https://github.com/takaram/kofam_scan). We annotated the mitochondrial genes with the GeSeq online tool (https://chlorobox.mpimp-golm.mpg.de/geseq.html). We annotated TEs with the Transposon Annotator “reasonaTE” tool (70) and calculated the percentage of the genome annotated as TEs by dividing the number of nucleotides in the annotated TE regions by the total genome length, multiplied by 100.

### Annotation of Aromatic C Degradation and N-acquisition Genes

We generated Hidden Markov Model (HMM) profiles for 18 aromatic C degradation and 4 N acquisition genes (Table S9) examined previously (27). Aromatic C degradation genes examined belonged to meta- and ortho-cleavage for both catechol and protocatechuate and naphthalene degradation pathways (26–28). According to previous studies, fungi transform aromatic C into intermediates such as catechol, protocatechuate, and benzoyl-CoA, followed by conversion to secondary intermediates such as acetyl-CoA, succinyl-CoA, and pyruvate through the β-ketoadipate (βKA) pathway (29), so we also examined genes from the bKA pathway.

We also examined N acquisition genes such as nitrate and nitrite reductase whose transcription were previously shown to increase after fire (71). We also included nitrous oxide reductase (*nosZ*) since high-severity fires may impair the soil microbiome’s capacity to convert nitrous oxide (N_2_O), a potent GHG, to nitrogen (N_2_) gas (72), leading to elevated N_2_O emission from burned soil (73). Fungi play a unique role in this process. We also included *P450nor* because some fungi possess Cytochrome P450 nitric oxide reductase (*P450nor*), an enzyme that catalyzes the reduction of nitric oxide (NO) to N_2_O during fungal denitrification (74).

We downloaded all protein sequences associated with each gene from NCBI, and discarded proteins annotated as similar or partial in their FASTA headers. We then aligned the high-confidence protein sequences of each gene using ClustalOmega v. 1.2.4 (75) with default parameters and used the alignments to build HMM profiles using the hmmbuild command of HMMER v.3.4 (http://hmmer.org/). We used these HMM profiles to annotate each fungal isolate’s protein sequences using the hmmsearch command of HMMER v.3.4, with an E-value cutoff of 1e-23. To confirm that the hmmsearch accurately retrieved the orthologues of each gene, we matched the available sequence descriptions and domain annotations obtained using BLAST and Interproscan with the description of proteins used as inputs for building HMM profiles.

For basidiomycetes with dikaryotic genomes, we assessed the contribution of internuclear rearrangements to the total number of aromatic C degradation and N acquisition genes by separately annotating the two haplotypes of genomes assembled with HiFiasm (58) using the FunGap pipeline (68). Coding sequences (CDSs) from the two haplotypes were then matched using JCVI (76). Gene relationships were categorized as 1:1 (one gene in each haplotype), duplicated (more than one copy in a single haplotype), or presence–absence variants (gene missing in one haplotype).

### Orthogroup Construction and Visualization

We performed comparative genomic analysis of 16 pyrophilous fungi belonging to the 6 orders for which publicly available genomes of sister species existed along with 38 sister species (Table S8) selected as described in Methods S3. We assigned genes to orthogroups (clusters of orthologues across genomes) using Orthofinder v.2.5.5 (77) and used the gene IDs of HMMER annotated aromatic C degradation and N acquisition genes to extract assigned orthogroups from the Phylogenetic Hierarchical Orthogroups predicted for the oldest common ancestor of all species (N0). We obtained the orthogroups per gene family and individual genes by creating a matrix of orthologue counts per species. We visualized the sum of genes for each of naphthalene degradation, catechol meta- and ortho-cleavage, protocatechuate (PCA) ortho- and meta-cleavage, and N-acquisition pathways alongside the species tree generated with Orthofinder using the plotTree.barplot function of Phytools (https://cran.r-project.org/web/packages/phytools/index.html) in R v.4.3.2 (78). To identify gene duplication events leading to the current composition of aromatic C degradation and N acquisition genes in the pyrophilous fungi, we obtained the number of gene duplication events in these genes occurring at each current node (53 species) and their common ancestors from the Orthofinder Gene Duplication Events reports. For dikaryotic genomes, we assessed the genome enrichment of each haplotype separately (supplementary file 1) and count of gene duplication events was doubled if both haplotypes carried the genes. We visualized the number of gene duplication events on the phylogenetic tree alongside the orthologue counts per individual gene using iTOL software v.7 (https://itol.embl.de/).

We also tested the involvement of phylogenetic signals in aromatic C and N acquisition gene enrichment. We normalized the gene counts by total proteome size to account for differences in genome content. Using the rooted phylogenetic tree generated by Orthofinder, we quantified phylogenetic signal for each gene with Pagel’s λ, estimating associated likelihood-ratio test p-values using the phylosig function from the R package phytools (79).

To test for enrichment of gene families in pyrophilous versus non-pyrophilous fungi, we implemented generalized linear models (GLMs) with a Poisson distribution in R (78), using gene family counts as the response variable, fungal lifestyle (pyrophilous or non-pyrophilous) as the predictor, and proteome size as an offset. Significance of enrichment was assessed from the trait (pyrophilous vs. non-pyrophilous) coefficient, and false discovery rate (FDR) was controlled using the Benjamini–Hochberg method.

### Identification of candidate HGT events

We identified candidates for HGT events by visually inspecting the putative xenologues generated by Orthofinder for outlier orthologues clustered distantly from their sister species in their gene trees then validated them with the following steps. We visualized the aligned protein sequences of candidate HGT genes and their corresponding native genes in the orthogroup using iTOL software v.7 (https://itol.embl.de/). Next, we compared candidate HGT gene transcription levels with their corresponding native genes (orthologues complying with phylogeny in the gene tree) in the orthogroup. We aligned the RNA-seq reads to their corresponding fungal genome using HISAT2 v.2.2.1 (80). We then calculated the Transcript Per Million (TPM) values for each gene using TPMCalculator v.0.0.5 (81) from the spliced alignment BAM file generated with HISAT2. Next, we calculated synonymous (S) and non-synonymous (NS) nucleotide substitution rates of the HGT candidates and their corresponding native genes using KaKs_Calculator v.3.0 (82). We also examined the co-localizations of the loci with the candidate HGT genes and TEs by visualizing the GFF3 files containing gene and TEs annotations in Integrated Genome Viewer (IGV) software v.2.15.2 (64). We also used BLASTP v.2.16.0 (https://blast.ncbi.nlm.nih.gov/Blast.cgi) to retrieve the protein sequences with homology to candidate HGT genes within all publicly available proteins from fungi and bacteria kingdoms in NCBI gene bank for phylogenetic analysis to identify potential HGT donor species. We included a *catA* gene identified on a conjugative plasmid of a pyrophilous bacteria *Noviherbaspirillum soli* (from Burkhorderiales order) through genome sequencing of pyrophilous bacteria (83).

### Measuring genetic variation among taxa in aromatic C degradation and N acquisition genes

We calculated the nucleotide substitution rate of the aromatic C degradation and N acquisition genes for the 53 genomes subjected to orthogroup construction with KaKs_Calculator v.3.0 (82). We visualized with a heatmap generated for a single orthogroup per each of the naphthalene degradation, catechol meta and ortho-cleavage, and protocatechuate meta and ortho-cleavage, and N acquisition pathways using the heatmap.2 function of R v.4.3.2 (84). The presence of phylogenetic signals and the significant differences between the pyrophilous and their non-pyrophilous sister species in the number of nucleotide substitutions were drawn as described above for gene enrichment analysis. See Methods S4 for methods examining contribution of internuclear variation in Basidiomycota to variation among aromatic C degradation and N acquisition genes.

### Measuring trade-off between fast-growth, and aromatic C degradation and N-acquisition

We measured HE rate on MYA or PyOM media for 16 filamentous pyrophilous isolates using three biological replicate plates per media type per isolate. We covered each plate with a sterile cellophane sheet (16), then inoculated a 5 mm² radius plugs of fungal tissue into the center, then incubated at room temperature with the hyphal growing edge recorded every 24 h using alternating-colored markers. After 3 weeks, plates were photographed, and images were imported into ImageJ v.1.54 (https://imagej.net/ij/) where daily extension rates were quantified in millimeters per day. We calculated the pairwise Spearman’s rank correlation between the HE rates on MYA and PyOM media and the number of each aromatic C degradation and N acquisition orthologues of each species using the rcorr function (85) and visualized using the corrplot function of R (86).

## Acknowledgements

This project was funded by the Department of Energy BER Award #DE-SC0023127 to SIG, PMH, SDA, and MJW and United States Department of Agriculture-NIFA Award #2022-67014-36675 to SIG and PMH. The RNA-seq studying transcription Post-PyOM treatment was conducted through Proposal ID: 510601, DOI: 10.46936/10.25585/60012549 by the U.S. Department of Energy Joint Genome Institute (https://ror.org/04xm1d337), a DOE Office of Science User Facility, supported by the Office of Science of the U.S. Department of Energy operated under Contract No. DE-AC02-05CH11231. We thank many individuals who contributed to the Glassman lab pyrophilous microbial culture including lab managers Judy Chung and James Randolph, lab assistants Ryan Quaal and Aishwarya Veerabahu, collaborators Tom Bruns and Monika Fischer, UCR PhD students Fabiola Pulido-Chavez, Arik Joukhajian, Esbeiry Cordova-Ortiz, and Mark Yacoub, and UCR undergraduates Anna Nguyen, Marely Vega, Jenna Maddox, Justin Diab, Wine-Jie Lipardo, Audrey Reichard, Priscilla Shultz, Jorge Ponce, and Nathan Heger. We thank Jason Stajich and his lab for advice and protocols for high molecular weight DNA extraction. We thank our anonymous reviewers for their helpful feedback on the manuscript.

