## Supplementary fig 1-12 for "Gene Duplication, Horizontal Gene Transfer, and Trait Trade-offs Drive Evolution of Post-Fire Resource Acquisition in Pyrophilous Fungi"

**Supplementary results**

**Results S1. Genome Assembly Statistics:** We believe that six out of 18 genomes sequenced in this study are completely novel reference genomes, which is supported by the absence of public genomes for *Anthracobia macrocystis*, *Rasamsonia columbiensis*, *Pholiota brunnescens*, and *Holtermanniella festucosa* and the phylogenomic analysis showing that *Alternaria sp.* forms a distinct branch within the described Pleosporales species (Fig. S1). Further, *Neurospora sp.* EDF-1 is a distinct species from *N. discreta* (Fig. S2A) and is phylogenetically distinct from 128 *Neurospora* species, including 42 strains of *N. discreta* (Fig. S2B).

The genome assembly of *Neurospora discreta* E-DF3 was highly contiguous (Table S3), with metrics comparable to the most recent reference genome, *N. discreta* FGSC8579 (<https://mycocosm.jgi.doe.gov/Neudi2/Neudi2.home.html>). Our genome assembly of *Coniochaeta hoffmannii* exhibited a 22-fold increase in N50 (3.2 Mb versus 148.5 Kb) compared to the existing reference genome (strain EXF-13287, GenBank accession GCA_030052785.1), with 4/10 contigs containing telomere sequences at both ends.

Finally, all 6 isolates of pyrophilous fungi previously sequenced using PacBio CLR technology (1, 2) showed dramatic reductions in number of contigs and increases in contig N50 values with higher percentage of complete BUSCO proteins in the PacBio HiFi assemblies for 4/6 isolates (Table S6). We substantially improved the *Basidioascus undulatus* genome assembly, reducing the number of contigs from 2,992 to 18 and achieving an 11-fold increase in contiguity (N50). Additionally, we provided the first gene annotation for this species, as the previous genome assembly lacked annotation.

We identified 2,900 heterozygous SNPs in *B. undulatus* , 39 in *H. festucosa*, 1,360,152 in *P. brunnescens*, and 1,072,632 in *L atratum*, suggesting that the two yeast basidiomycetes *B.* *undulatus* and *H. festucosa* has haploid genome or has dikaryotic haploid genome with subtle internuclear genetic variation (Table S4). We found that 98% and 99% percent of internuclear SNPs were phased into two distinct haplotypes in *P. brunnescens* and *L. atratum*, supporting significant internuclear variation. The phased block N50 were 657 and 870 kb for *P. brunnescens* and *L. atratum*, respectively.

**Results S2. Variation in the Enrichment of Aromatic C Degradation and N Acquisition Orthologues Among Pyrophilous Fungi**: We performed comparative genomic analysis by assigning genes to orthogroups (clusters of orthologues across genomes) using Orthofinder software. To improve the confidence of orthogroup assignment, we added 38 public genomes of sister species to the analyses (Table S7). We could not include *Basidioascus undulatus* from Geminibasidiales and *Holtermanniella festucosa* from Holtermanniales in this analysis since they were the only genomes for these orders. The number of orthologues related to 18 aromatic C degradation and four N acquisition genes (Table S8) varied among the 53 taxa and six orders used to construct orthogroups (Fig. S3). Eurotiales had the most genes for aromatic C degradation, followed by Pleosporales, then Coniochaetales and the lowest in the Sordariales, Pezizales and Agaricales. While variation was less than that of the aromatic C degradation pathways, nonetheless N acquisition pathways exhibited variation among orders with the most N-acquisition gene orthologues in Coniochaetales and the fewest in Pezizales (Fig. S3).

While groupings of orders emerged indicating phylogenetic signatures, there was also variation within individual taxa within the orders in the number of orthologues for each of the aromatic C degradation pathways. For example, although the species belonging to the Pezizales order under Pyronemataceae family i.e., three pyrophilous species *Anthracobia macrocystis*, *Tricharina praecox* and *Pyronema omphalodes* had some of the lowest enrichment of catechol ortho-cleavage genes, the three *Morchella* species had the enrichment comparable with some Eurotiales (Fig. S3). Further, some pyrophilous species exhibited distinctively more aromatic C degradation genes than their sister isolates. For example, *Coniochaeta hoffmannii* had more orthologues for aromatic C degradation (particularly genes for naphthalene degradation and catechol meta-cleavage) than its sister isolate *C. hoffmannii* EXF-13287, which was isolated from a car fuel reservoir (3). *Lyophyllum atratum* also had more aromatic C degradation genes than its sister species *L. shimeji* AT787, an ectomycorrhizal fungus, due to having a higher number of bKA pathway orthologues. However, this same pattern was not observed for N acquisition genes.

Finally, our analyses suggest that the expansion in aromatic C degradation and N acquisition orthologues is not associated with expansion of genome sizes, but rather by selection for those genes within pyrophilous taxa (Fig. S3). For example, the largest genomes belonged to *Pyronema omphalodes* and *Peziza echinospora* from the Pezizales order, which had the lowest enrichment of aromatic C degradation and N acquisition orthologues.

**Supplementary methods**

**Method S1. HMW DNA extraction:** We grew fungi in Malt Yeast Broth (MYB) (Malt extract [RPI Research International, Illinois, USA] 5g, Yeast extract [Apex Chemical, Singapore] 5g, distilled water 1L) at room temperature (RT) for 5–10 days based on growth rate. For 7 isolates, we separated mycelia by pouring the entire flask over a funnel covered with sterile Whatman paper, washed the mycelia twice with sterile deionized water, dried by blotting between two sterile Whatman papers, then ground tissue in liquid N with mortar and pestle. Due to slow growth in MYB media, we grew *Peziza echinospora* and *Geopyxis carbonaria* on sterile cellophane sheets laid over Malt Yeast Agar (MYA)(MYB + 15 g agar [Apex Chemicals, Singapore]) plates for 10 days prior to scraping mycelia with a sterile scalpel and grinding in liquid N. For three yeast species, *Coniochaeta hoffmannii* (dimorphic, growing as a yeast in MYB), *Holtermanniella festucosa*, and *Basidioascus undulatus*, the cells were pelleted by centrifuging at 3000 rpm for 3 minutes. We extracted HMW DNA from fungal tissues with Qiagen Genomic Tip 100/G.

We extracted HMW DNA for 6 isolates (*Aspergillus cf. fellis*, *Penicillium antarcticum*, *Tricharina praecox*, *Pyronema omphalodes*, *Neurospora sp.* E-DF1, and *Neurospora discreta* E-DF3) using a CTAB:Phenol:Chloroform method (4) since extraction with Qiagen Tips led to low DNA concentration (Table S1).

We evaluated DNA purity and concentration with the Qubit™ Fluorometer dsDNA BR Assay Kit (Invitrogen, Carlsbad, USA) using a DeNovix spectrophotometer Model Ds-11 (DeNovix Inc. Wilmington, USA). We measured DNA integrity and fragment size with the Genomic DNA Screen Tape (Agilent, Waldbronn, Germany) at the University of California, Riverside Institute for Integrative Genome Biology (UCR IIGB).

**Methods S2. RNA extraction and quality control:** RNA was extracted from fungal isolates grown under three distinct conditions as detailed in the main text of the paper. We collected mycelia from the liquid media using the same methods for HMW DNA extraction. We pooled mycelia from all three conditions (described in the main text) and ground in a mortar and pestle chilled with liquid N. We assessed RNA purity and concentration by a Qubit™ RNA Broad Range Assay Kit (Invitrogen, Carlsbad, USA) using a DeNovix spectrophotometer Model Ds-11 (DeNovix Inc. Wilmington, USA) and determined RNA integrity using an Agilent 2100 Bioanalyzer (Agilent Technologies Inc., Santa Clara, USA).

**Methods S3. Selection of sister species for comparative genomic analysis:** For the 6 orders included for comparative genomic analysis, we downloaded the protein FASTA files of all publicly available genomes using the get_assembly tool v.0.10.0 and by specifying the order name in the “organism” option (<https://github.com/davised/get_assemblies>). We then constructed a phylogenomic tree for each individual order using the BUSCO_phylogenomics tool release 20240919 (<https://github.com/jamiemcg/BUSCO_phylogenomics>). We used the protein FASTA file of each genome as input to conduct BUSCO analysis (5) and obtain the sequence of shared BUSCO proteins for multiple sequence alignment. We constructed the phylogenomic tree from the trimmed alignment generated with BUSCO_phylogenomics using FastTree v2.1.11 (6) software with -spr 4, -mlacc 2, -slownni, and -gamma parameters. We assigned the sister species to be those most closely clustered with the target genomes.

**Methods S4. The contribution of inter-nuclear variation in basidiomycetes to N-acquisition genes:** The basidiomycota species exhibited the highest synonymous and non-synonymous substitutions in an orthogroup annotated as nitrate reductase (*narG-narZ*). We evaluated the contribution of inter-nuclear variation to this higher nucleotide polymorphism in *Lyophyllum atratum* as an example. We retrieved two haplotypes derived from two nuclei of a gene annotated as *narG-narZ* in *Lyophyllum atratum* (LA_1440.t1) from the phased genome assembly using BLASTn v.2.16.0 (<https://blast.ncbi.nlm.nih.gov/doc/blast-help/downloadblastdata.html#downloadblastdata>) and aligned the retrieved cDNA sequences using minimap2 v.2.18 (7). We used the DNA sequence of one haplotype as a reference for aligning the RNA-seq reads using HISAT2 v.2.2.1 (8). We imported the BAM files containing the alignments into IGV software v.2.15.2 (9) to visualize the genetic variation between the haplotypes and the contribution of each to the total transcription levels of the *narG-narZ*.

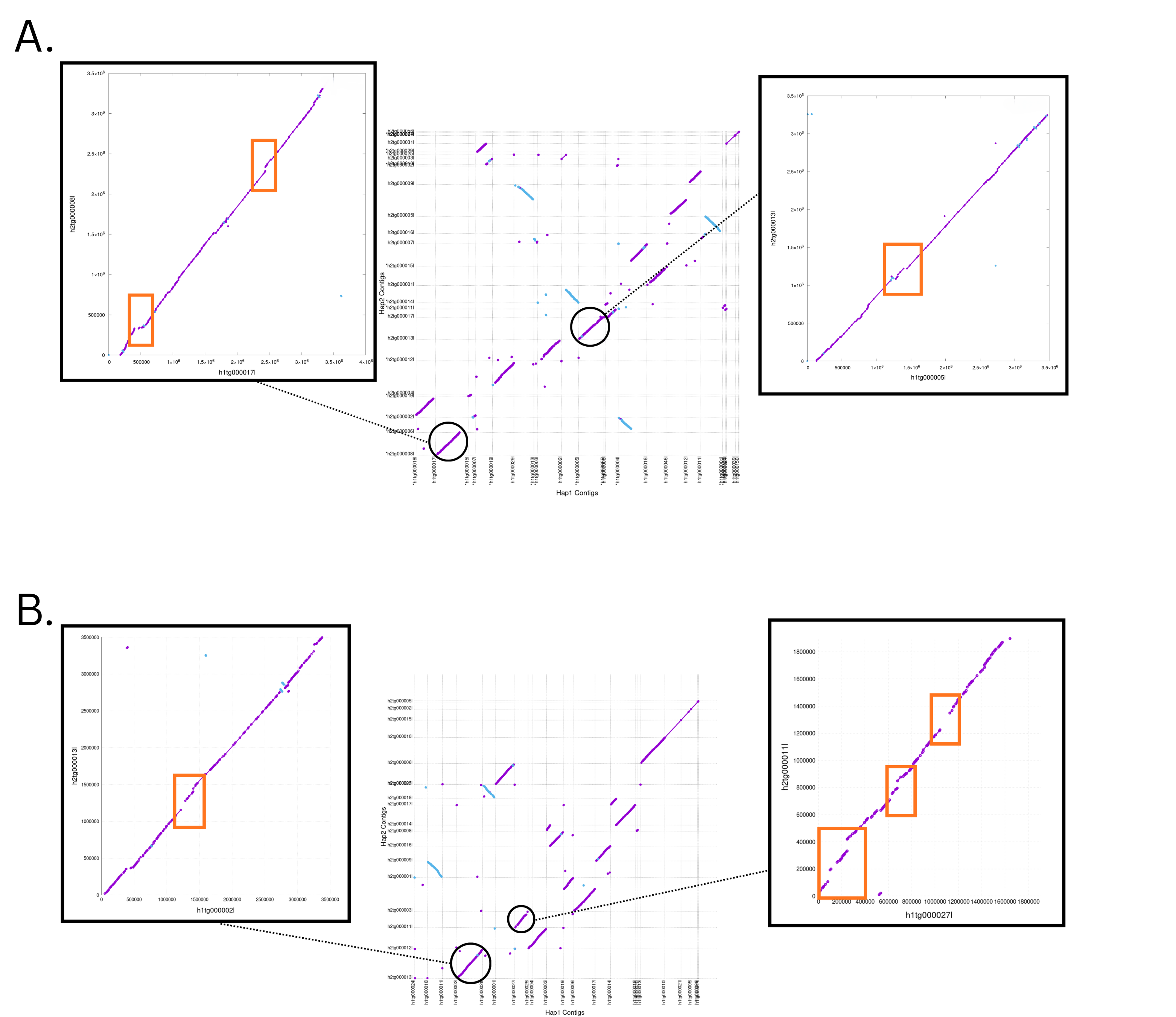

**Fig S1.** The sequence alignment between the contigs representing two haplotypes of *Pholiota brunnescens* (A) and *Lyophyllum atratum* (B) genomes. Forward alignments are shown as purple lines and reverse alignments as blue lines. The central plot displays alignments of all contigs, while the side panels provide zoomed-in views of two example contig alignments, with rearrangements highlighted by orange rectangles.

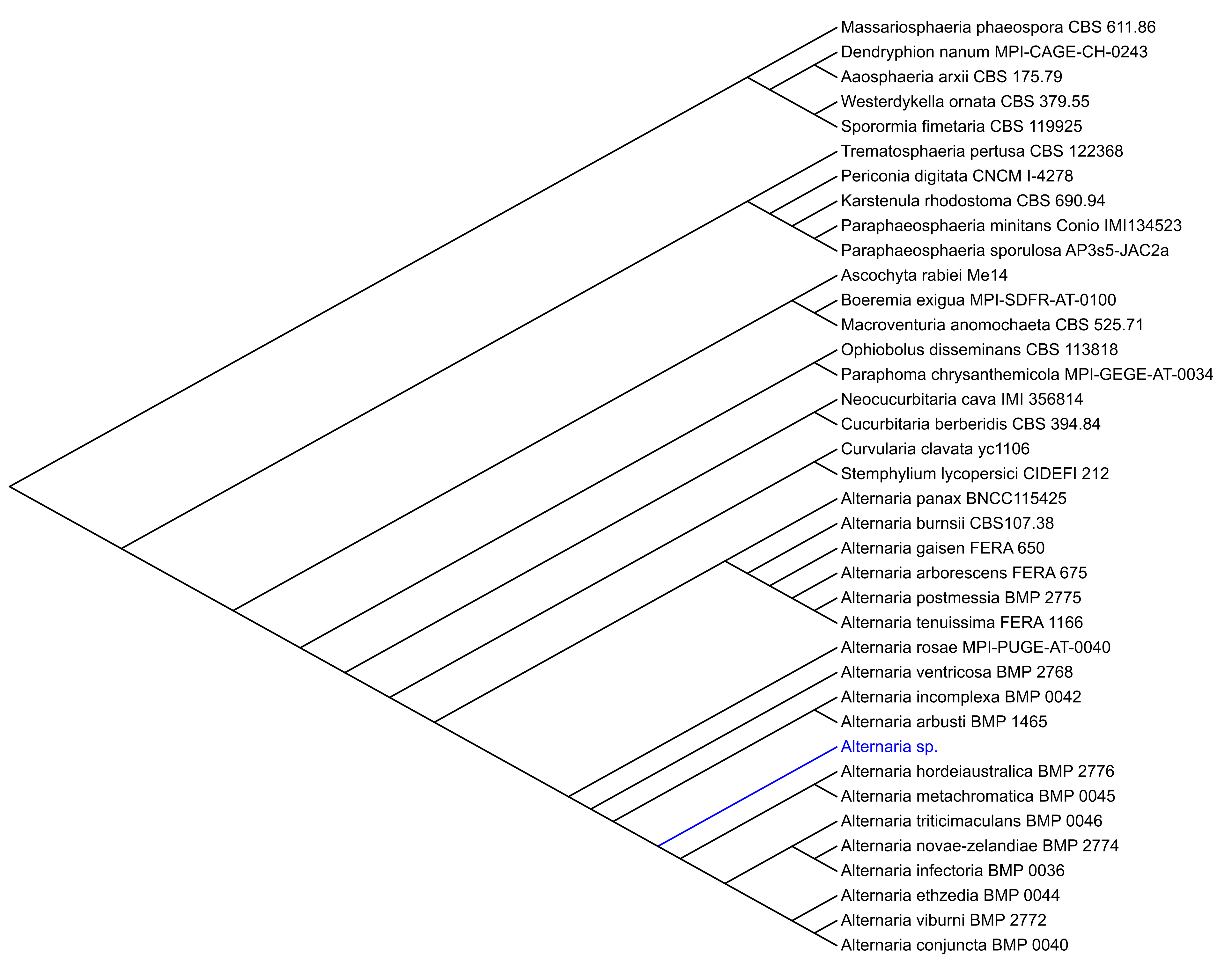

**Fig S2.** Phylogenomic tree of all the Pleosporales species with public genomes in NCBI vs. *Alternaria* sp. sequenced in this project (blue ink).

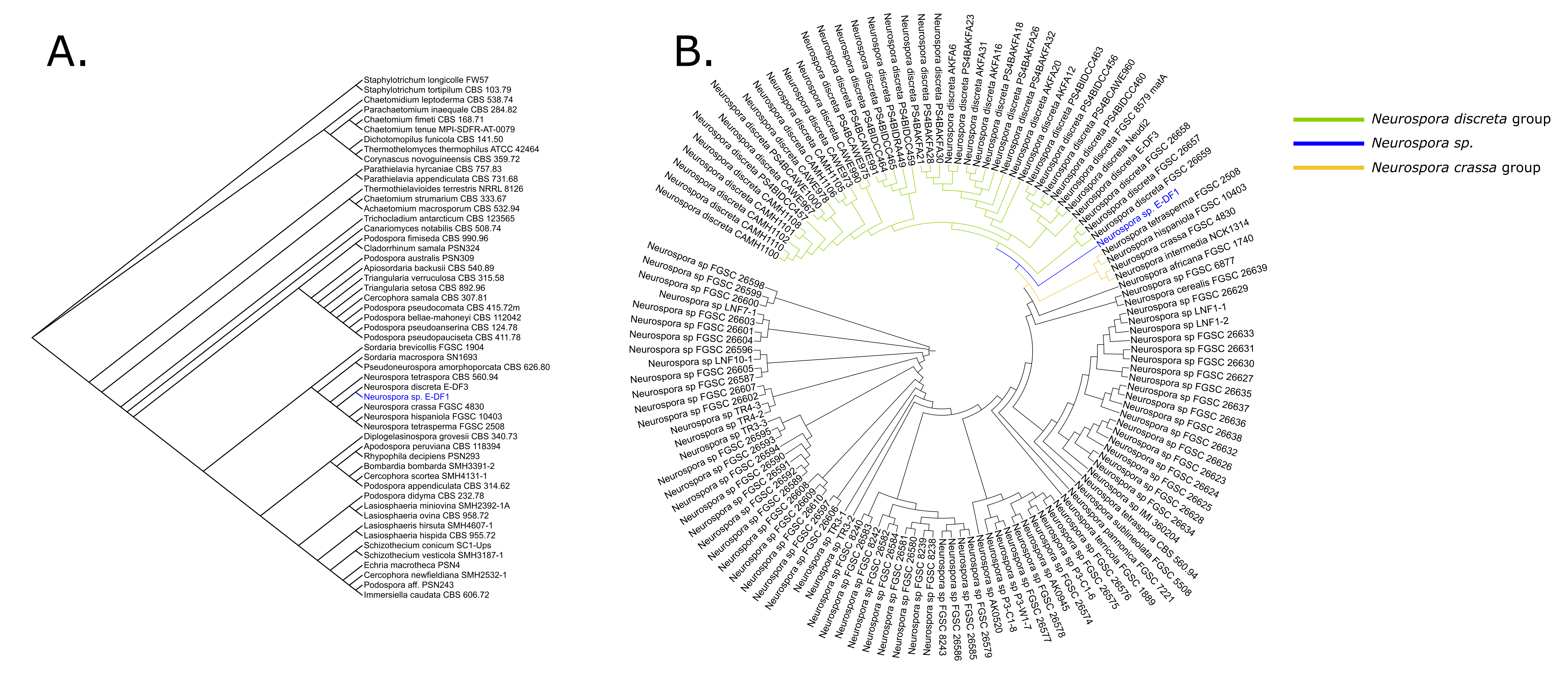
**Fig S3.** Phylogenomic tree of all the Sordariales species with public genomes and *Neurospora* sp. E-DF1 sequenced in this study (blue ink) **(A)** and phylogenomic tree of all the Neurospora species with public genomes along with *N.* sp. E-DF1 (blue ink) **(B)**. The green branches are all *N. discreta* isolates and orange branch contains *N. crassa* and some phylogenetically related species

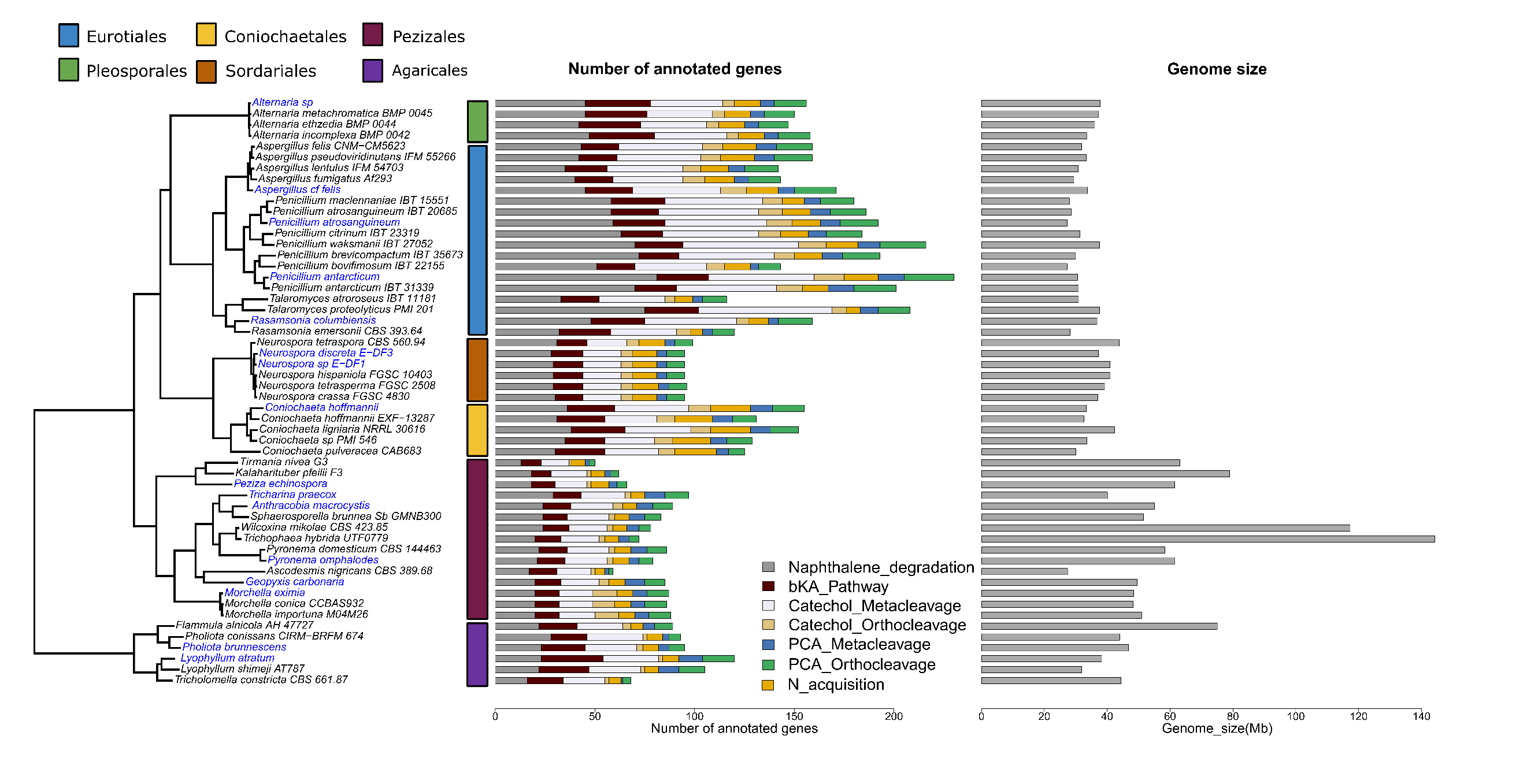
**Fig S4.** Number of annotated aromatic carbon (C) degradation (six pathways) and nitrogen (N) acquisition orthologues vs. the genome size of 53 genomes used for constructing orthogroups across 6 orders of pyrophilous fungi. The tree leaves in blue text are 16 pyrophilous species sequenced in this study.

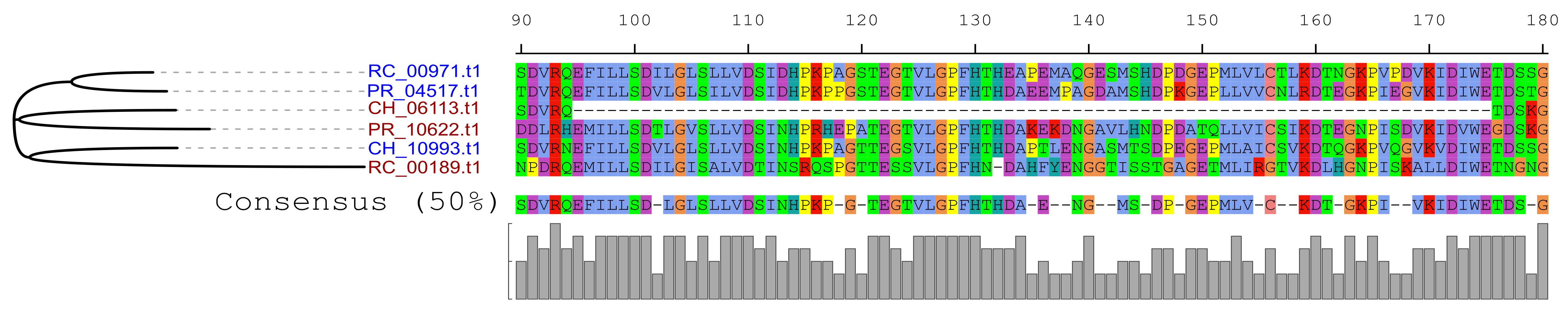

**Fig S5.** The protein sequence alignment of three candidates horizontally transferred genes (brown tree leaves) in *Coniochaeta hoffmannii* (CH_06113.t1), *Penicillium atrosanguineum* (PR_10622.t1), and *Rasamsonia columbiensis* (RC_00189.t1) along with their native paralogues (blue tree leaves) within a catechol-1,2-dioxygenase (*catA*) encoding orthogroup.

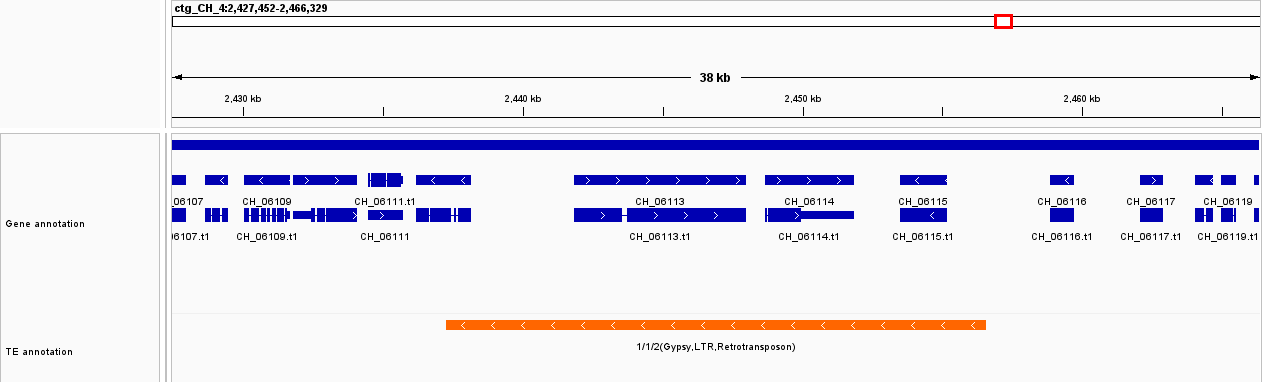

**Fig S6.** The co-localization of a Ty3-like (Gypsy) Long Terminal Repeat (LTR) retrotransposons with CH_06113.t1, a candidate horizontally transferred gene in *Coniochaeta hoffmannii.* Top panel shows the genome coordinate of the contig, and the red box is the location of the gene on the contig. The bottom panel presents the position of genes (blue bars) and the transposable element (red bar).

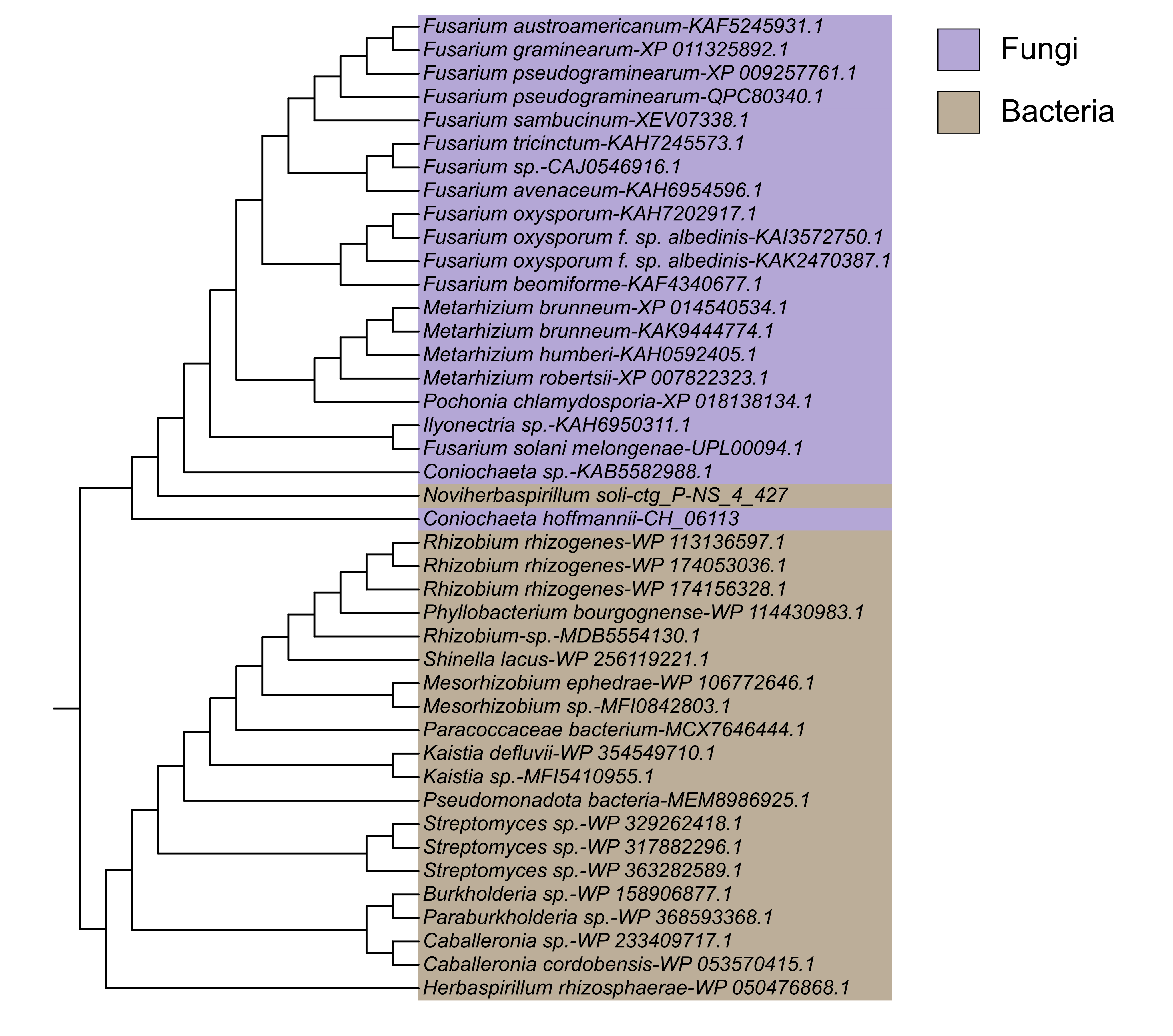

**Fig S7.** The phylogenetic tree of the CH_06113.t1, a candidate horizontally transferred gene in *Coniochaeta hoffmannii,* and its orthologues in other fungi (Blue shade) and bacteria (Red shade) obtained through NCBI BLASTP search, and an orthologue (*ctg_P-NS_4_427*) we found on a conjugative plasmid of *Noviherbaspirillum soli*, a pyrophilous bacteria isolated from the same habitat. The tree leaves are named with the species name followed by the Uniprot ID of the BLASTP hits.

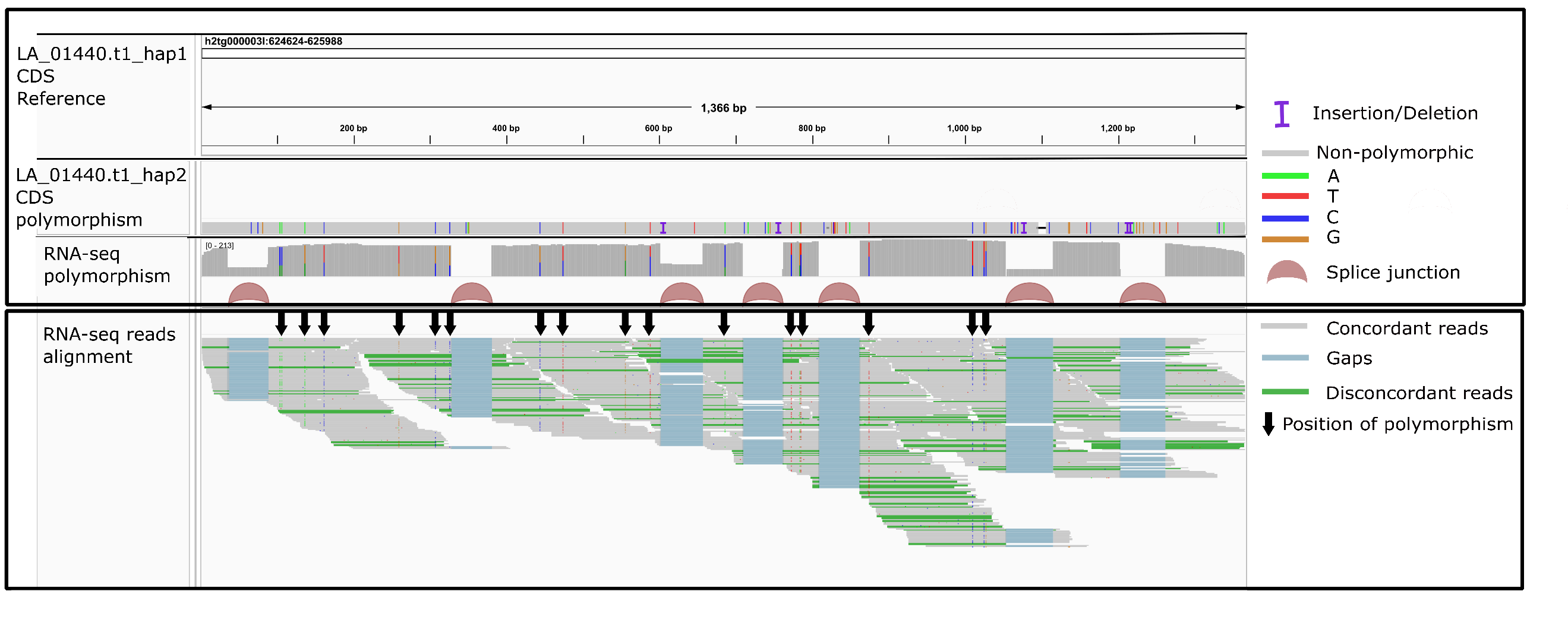

**Fig S8.** Internuclear variation contributing to nitrate reductase (*narG-narZ*) gene (LA_01440.t1) variation in dikaryotic basidiomycete pyrophilous fungi *Lyophyllum atratum*. The coding sequence (CDS) of two haplotypes of LA_01440.t1 gene (hap1 and hap2) were aligned to identify polymorphism sites (shown with black arrows). The top panel shows the snapshot of alignment of two haplotypes and the RNA-seq reads alignment to LA_01440.t1 haplotype 1 (LA_01440.t1_hap1). The bottom panel shows the RNA-seq reads and their alignment highlighting the presence of both haplotypes in the transcriptome.

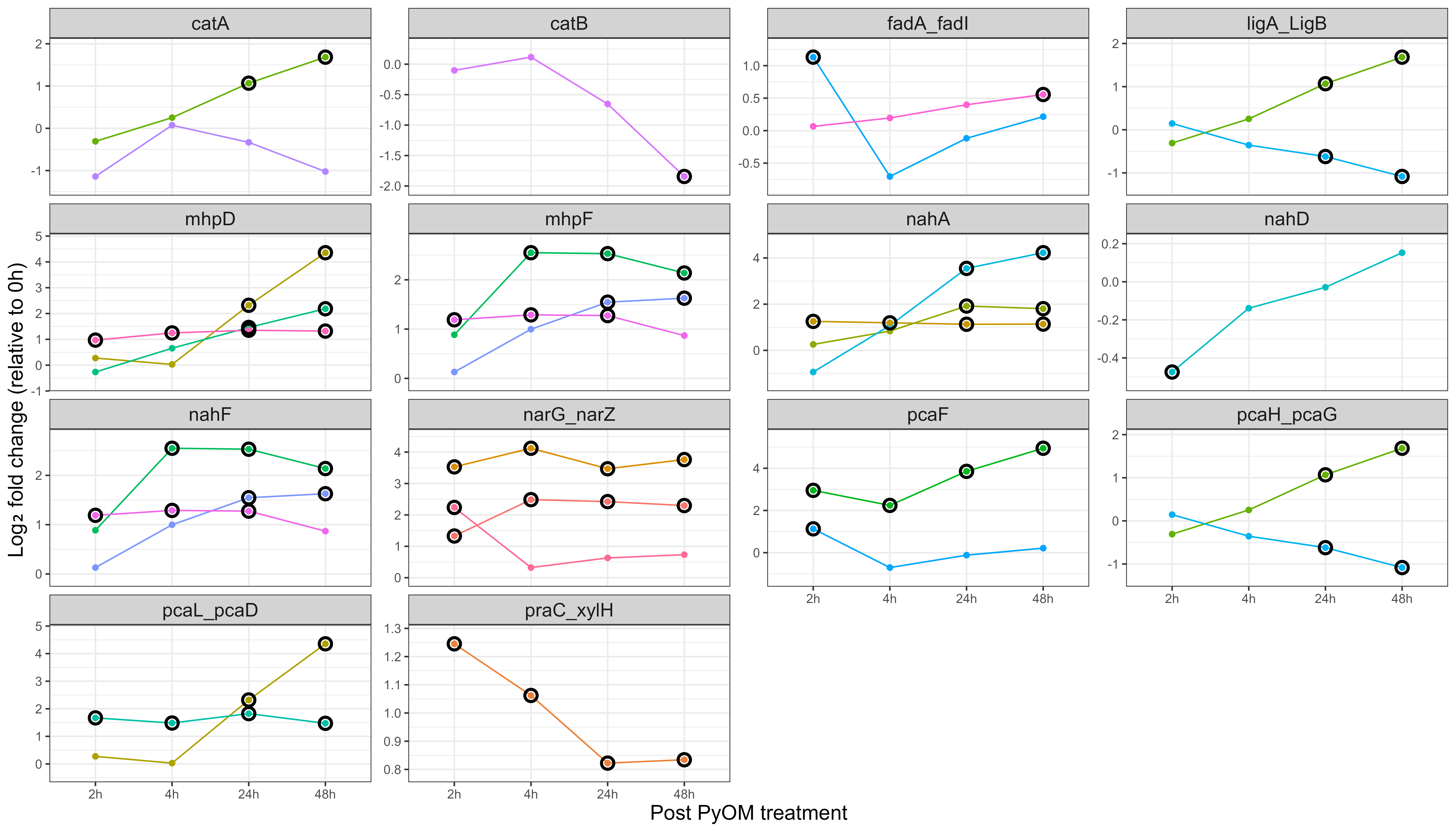

**Fig. S9.** Transcriptional changes of selected aromatic C and N acquisition genes at 2, 4, 24, and 48 h after PyOM treatment in the pyrophilous fungus *Coniochaeta hoffmannii*. Log_2_ fold changes were calculated relative to the 0 h time point (immediately before PyOM treatment). Statistical significance of differential transcription was determined using adjusted p-values (Benjamini–Hochberg false discovery rate < 0.05) and is indicated by black circles around data points in the line plots. Multiple line plots in each panel represent different orthologues of the same gene. Gene symbols are those in Table S9.

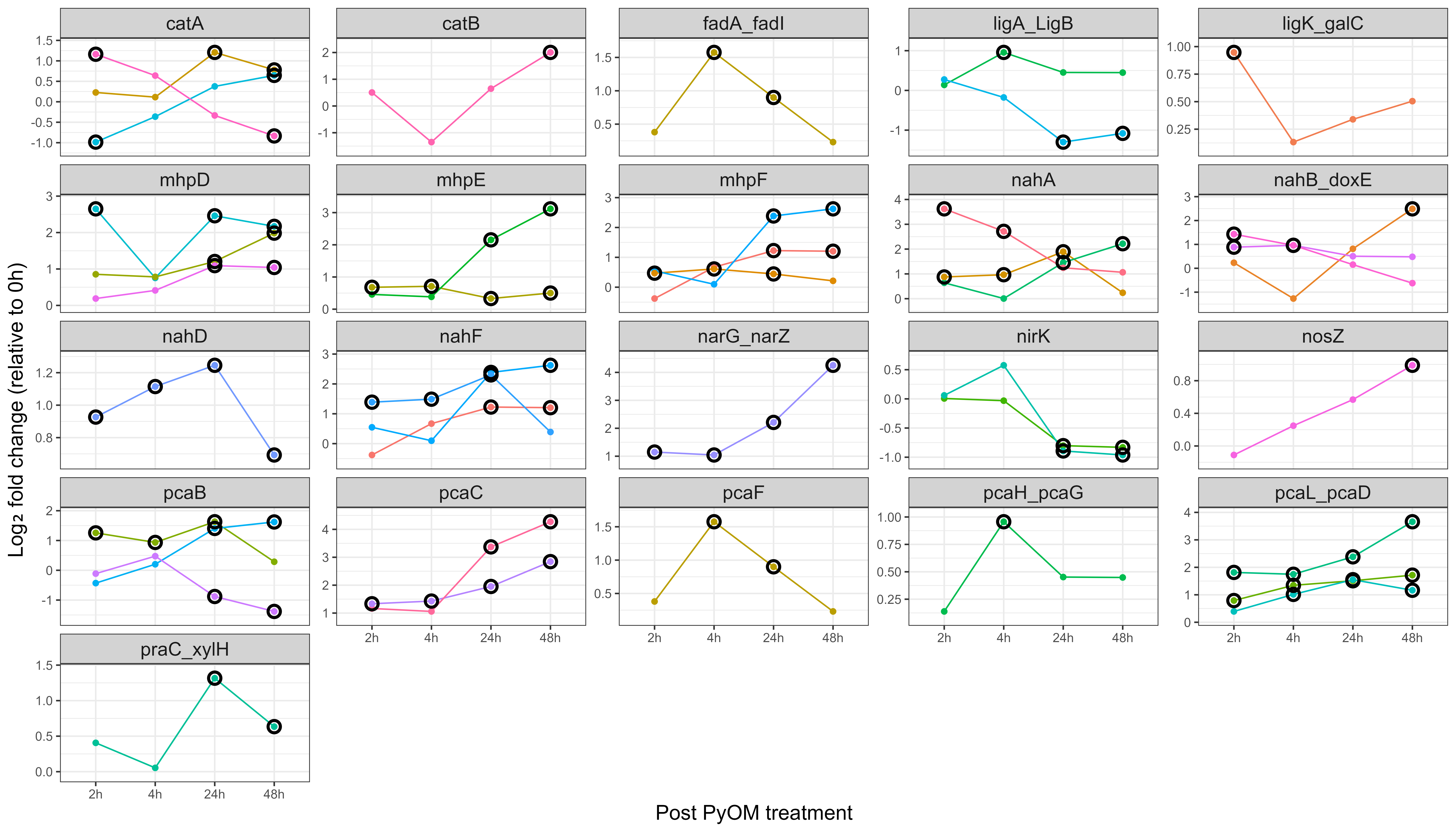

**Fig. S10.** Transcriptional changes of selected aromatic C and N acquisition genes at 2, 4, 24, and 48 h after PyOM treatment in the pyrophilous fungus *Penicillium atrosanguineum*. Log_2_ fold changes were calculated relative to the 0 h time point (immediately before PyOM treatment). Statistical significance of differential transcription was determined using adjusted p-values (Benjamini–Hochberg false discovery rate < 0.05) and is indicated by black circles around data points in the line plots. Multiple line plots in each panel represent different orthologues of the same gene. Gene symbols are those in Table S9.

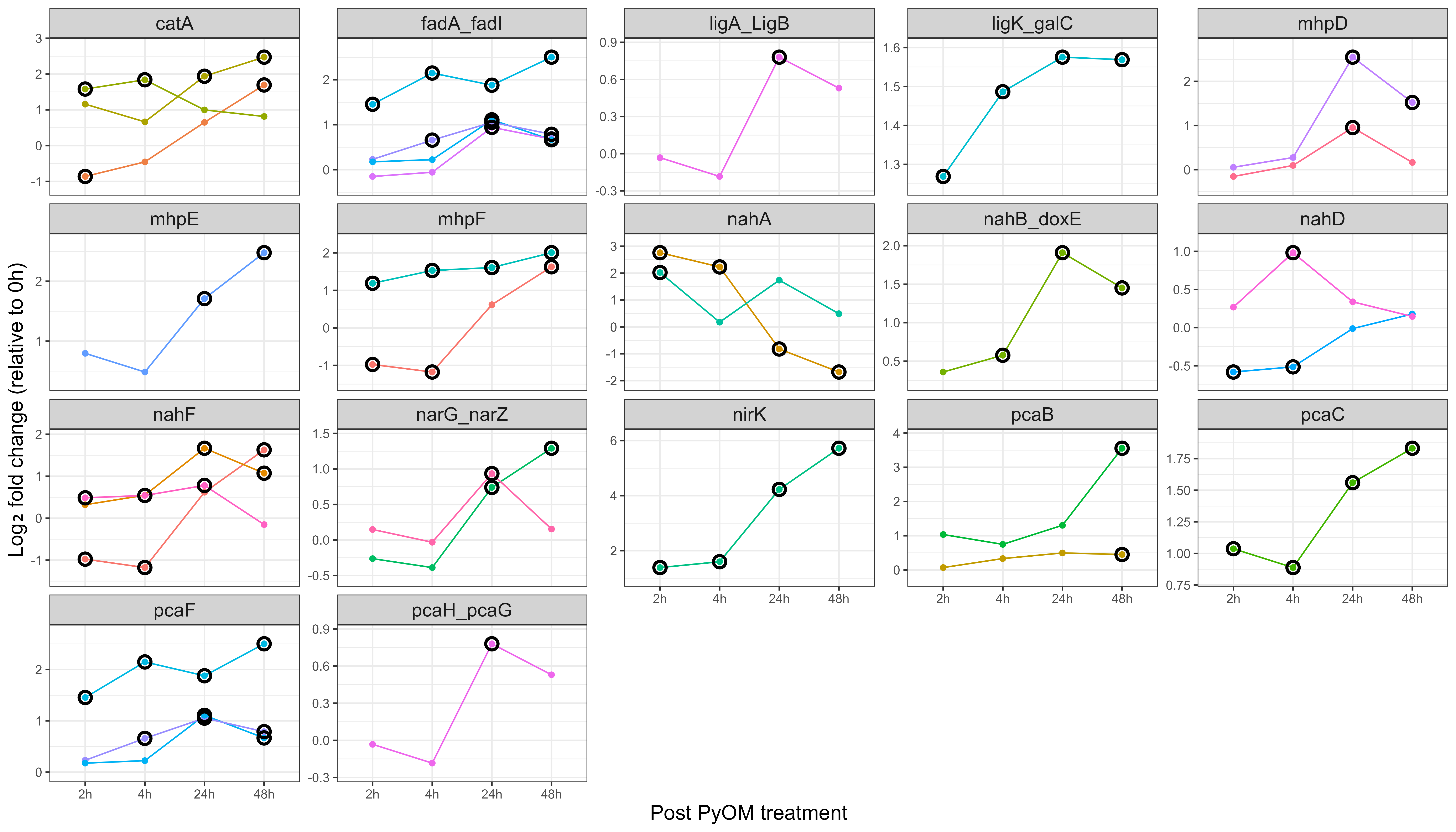

**Fig. S11.** Transcriptional changes of selected aromatic C and N acquisition genes at 2, 4, 24, and 48 h after PyOM treatment in the pyrophilous fungus *Morchella eximia*. Log_2_ fold changes were calculated relative to the 0 h time point (immediately before PyOM treatment). Statistical significance of differential transcription was determined using adjusted p-values (Benjamini–Hochberg false discovery rate < 0.05) and is indicated by black circles around data points in the line plots. Multiple line plots in each panel represent different orthologues of the same gene. Gene symbols are those in Table S9.

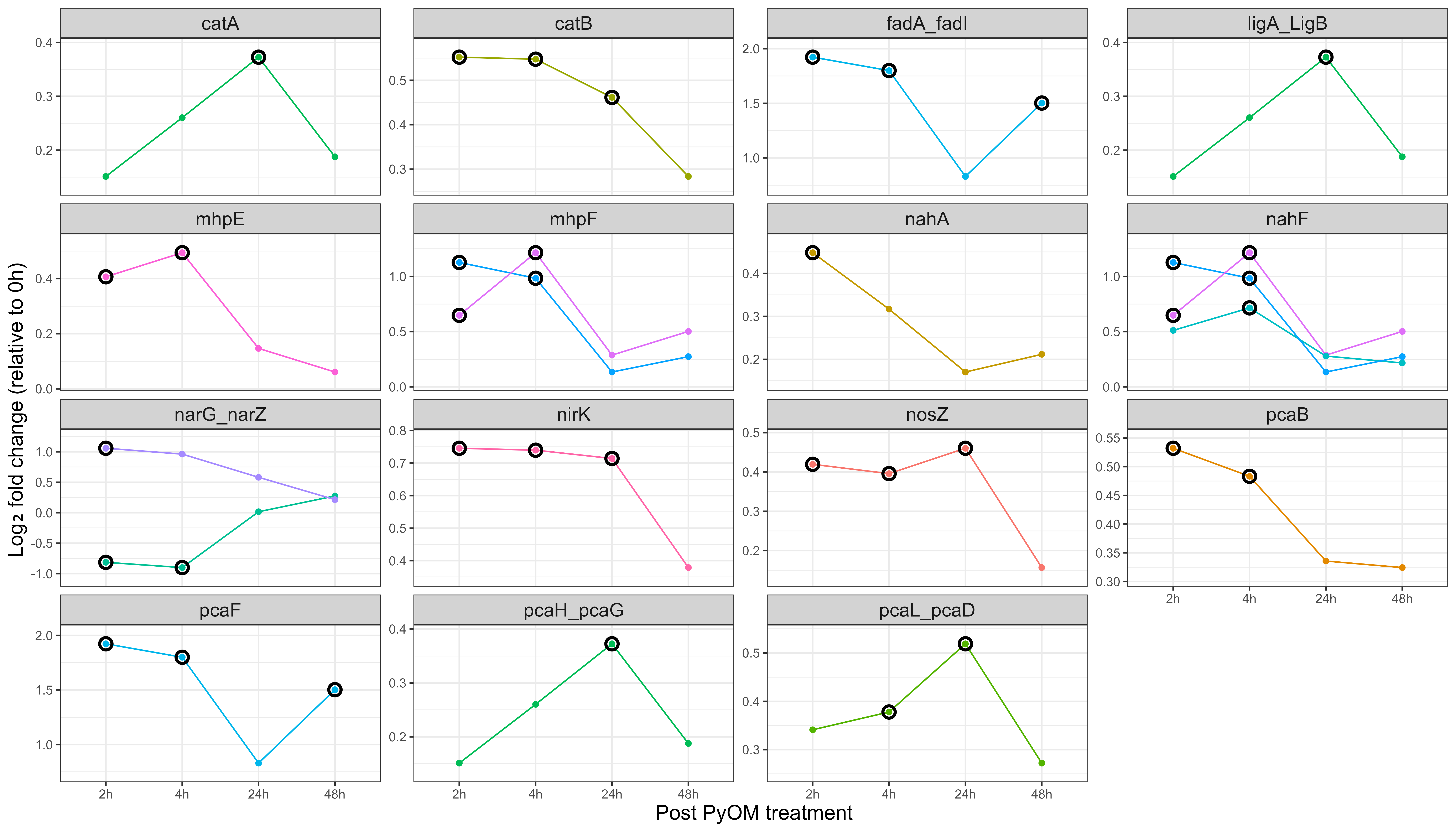

**Fig. S12.** Transcriptional changes of selected aromatic C and N acquisition genes at 2, 4, 24, and 48 h after PyOM treatment in the pyrophilous fungus *Pholiota brunnescens*. Log_2_ fold changes were calculated relative to the 0 h time point (immediately before PyOM treatment). Statistical significance of differential transcription was determined using adjusted p-values (Benjamini–Hochberg false discovery rate < 0.05) and is indicated by black circles around data points in the line plots. Multiple line plots in each panel represent different orthologues of the same gene. Gene symbols are those in Table S9.

**Table S1.** The 18 fungal isolates whose genomes were assembled in this study. All our fungi were isolated from burned soils or mushrooms collected after four California wildfires except for *Basidioascus undulatus* DAOM241956, which is a heat-resistant isolate from Canadian soil obtained from the Canadian National Mycological Herbarium. High molecular weight DNA was extracted for all species using the Qiagen Genomic Tip kit except for those with * extracted using a CTAB:Phenol:Chloroform method. Media types: Malt Yeast extract Agar (MYA), Pyrogenic Organic Matter (PyOM), Reasoner’s 2 Agar (R2A), Cornmeal Yeast Malt Agar (CMYM), Malt extract Agar (MA), Potato Dextrose Agar (PDA).

| **Phylum** | **Order** | **Species** | **Isolation source** | **Isolate ID** | **Wildfire/year/county/State** | **Isolation media** |
| --- | --- | --- | --- | --- | --- | --- |
| Ascomycota | Eurotiales | *Penicillium atrosanguineum* | soil | PYOM_F_13 | El Dorado Fire, El Dorado Ranch Park, 2020, San Bernardino County, CA | PyOM |
|  | Eurotiales | *Penicillium antarcticum** | soil | F-501 | Sycamore Canyon Fire, Riverside, 2019, Riverside County, CA | MYA |
|  | Eurotiales | *Aspergilus cf. fellis** | soil | T4_F18 | El Dorado Fire, El Dorado Ranch Park, 2020, San Bernardino County, CA | MYA |
|  | Eurotiales | *Rasamsonia* *columbiensis* | soil | R2A_F_01 | El Dorado Fire, El Dorado Ranch Park, 2020, San Bernardino County, CA | R2A |
|  | Pleosporales | *Alternaria sp.* | soil | PSF_3 | Sycamore Canyon Fire, Riverside, 2019, Riverside County, CA | MYA |
|  | Coniochaetales | *Coniochaeta hoffmannii* | soil | R2A_F_24 | El Dorado Fire, El Dorado Ranch Park, 2020, San Bernardino County, CA | R2A |
|  | Sordariales | *Neurospora sp.** | Joshua tree burnt bark | E-DF1 | Dome Fire, Mojave National Preserve, 2020, San Bernardino County, CA | MYA |
|  | Sordariales | *Neurospora discreta** | Joshua tree burnt bark | E-DF3 | Dome Fire, Mojave National Preserve, 2020, San Bernardino County, CA | MYA |
|  | Pezizales | *Peziza echinospora* | Apothecia | CBS144458 | Rim Fire, Stanislaus National Forest, 2013, Tuolumne County, CA | CMYM |
|  |  | *Anthracobia macrocystis* | Apothecia | N/A | Rim Fire, Stanislaus National Forest, 2013, Tuolumne County, CA | CMYM |
|  |  | *Geopyxis carbonaria* | Apothecia | CBS144460 | Rim Fire, Stanislaus National Forest, 2013, Tuolumne County, CA | CMYM |
|  |  | *Pyronema omphalodes** | Apothecia | CBS144459 | Rim Fire, Stanislaus National Forest, 2013, Tuolumne County, CA | CMYM |
|  |  | *Morchella eximia* | Apothecia | CBS144464 | Rim Fire, Stanislaus National Forest, 2013, Tuolumne County, CA | CMYM |
|  |  | *Tricharina praecox** | Apothecia | CBS144465 | Rim Fire, Stanislaus National Forest, 2013, Tuolumne County, CA | CMYM |
| Basidiomycota | Agaricales | *Pholiota brunnescens* | Mushroom | EDT7_PBRUN | El Dorado Fire, El Dorado Ranch Park, 2020, San Bernardino and Riverside Counties, CA | MYA |
|  |  | *Lyophyllum atratum* | Mushroom | CBS14462 | Rim Fire, Stanislaus National Forest, 2013, Tuolumne County, CA | MA |
|  | Holtermanniales | *Holtermanniella festucosa* | Soil | R2A_Y_01 | El Dorado Fire, El Dorado Ranch Park, 2020, San Bernardino County, CA | R2A |
|  | Geminibasidiales | *Basidioascus undulatus* | soil | DAOM241956 | - | PDA |

**Table S2.** Summary of PacBio HiFi sequencing data for 18 pyrophilous fungi, including total base pair (bp) sequence generated, read N50, and estimated genome coverage

| Species | Total base (Gbp) | Read N50 (bp) | Genome coverage (X) |
| --- | --- | --- | --- |
| *Penicillium atrosanguineum* | 18.8 | 6,478 | 59 |
| *Penicillium antarcticum* | 26.9 | 10,028 | 83 |
| *Aspergilus cf. fellis* | 27.2 | 8,654 | 94 |
| *Rasamsonia columbiensis* | 20.2 | 7,762 | 72 |
| *Alternaria sp.* | 24.6 | 7,545 | 77 |
| *Coniochaeta hoffmannii* | 6.5 | 6,418 | 21 |
| *Neurospora sp.* | 106 | 7,848 | 258 |
| *Neurospora discreta* | 213.7 | 7,323 | 521 |
| *Peziza echinospora* | 33 | 6,478 | 52 |
| *Anthracobia macrocystis* | 23 | 6,926 | 45 |
| *Geopyxis carbonaria* | 28.2 | 5,429 | 56 |
| *Pyronema omphalodes* | 51.8 | 6,030 | 85 |
| *Morchella eximia* | 186.5 | 6,269 | 252 |
| *Tricharina praecox* | 119.1 | 7,389 | 305 |
| *Pholiota brunnescens* | 33.2 | 7,708 | 25 |
| *Lyophyllum atratum* | 35.8 | 7,447 | 28 |
| *Holtermanniella festucosa* | 58.2 | 8,857 | 58 |
| *Basidioascus undulatus* | 21 | 8,633 | 33 |

**Table S3.** The genome assembly statistics i.e., genome size, contiguity, ploidy, and Transposable Elements (TEs) contents and NCBI project assigned to each genome. * Pyrophilous fungal isolates sequenced for the first time in this project, † those sequenced previously using PacBio CLR technology or Illumina short reads, with improved assembly metrics, and # those with no prior reference genome. For dikaryotic haploid assemblies, statistics correspond to the primary haplotype representative.

| **Species** | **# contigs** | **Total length (Mbp)** | **N50 (Mbp)** | **L50** | **TEs (%)** | **Genome ploidy** | **NCBI project** |
| --- | --- | --- | --- | --- | --- | --- | --- |
| *Penicillium atrosanguineum** | 8 | 27.3 | 3.9 | 4 | 6 | Haploid | PRJNA1251102 |
| *Penicillium antarcticum** | 8 | 30.6 | 4.8 | 3 | 8 | Haploid | PRJNA1251081 |
| *Aspergilus* cf. *fellis** | 10 | 33.7 | 4.2 | 4 | 9 | Haploid | PRJNA1249956 |
| *Rasamsonia columbiensis*# | 14 | 36.7 | 5.4 | 3 | 11 | Haploid | PRJNA1251120 |
| *Peziza echinospora*† | 38 | 61.4 | 2.5 | 10 | 19 | Haploid | PRJNA1251112 |
| *Anthracobia macrocystis*# | 33 | 55 | 2.3 | 7 | 25 | Haploid | PRJNA1247549 |
| *Geopyxis carbonaria*† | 35 | 49.5 | 1.6 | 13 | 19 | Haploid | PRJNA1250168 |
| *Pyronema omphalodes*† | 46 | 61.6 | 2.1 | 10 | 24 | Haploid | PRJNA1251119 |
| *Morchella eximia*† | 42 | 48.4 | 1.5 | 13 | 13 | Haploid | PRJNA1250785 |
| *Tricharina praecox*† | 41 | 40 | 1.8 | 8 | 9 | Haploid | PRJNA1251124 |
| *Alternaria* sp*.*# | 32 | 37.8 | 2.5 | 5 | 10 | Haploid | PRJNA1248598 |
| *Coniochaeta hoffmannii** | 10 | 33.5 | 3.2 | 5 | 4 | Haploid | PRJNA1250163 |
| *Neurospora* sp. E-DF1# | 23 | 41 | 1.4 | 10 | 14 | Haploid | PRJNA1251080 |
| *Neurospora discreta* E-DF3*** | 7 | 37.2 | 9.2 | 2 | 6 | Haploid | PRJNA1251063 |
| *Holtermanniella festucosa*# | 14 | 16.6 | 1.1 | 6 | 5 | Haploid | PRJNA1250175 |
| *Basidioascus undulatus*† | 18 | 34.8 | 2.2 | 6 | 7 | Haploid | PRJNA1250154 |
| *Lyophyllum atratum*† | 27 | 38.2 | 2.6 | 6 | 18 | Haploid dikaryotic | PRJNA1250761 |
| *Pholiota brunnescens*# | 49 | 46.8 | 3.8 | 6 | 22 | Haploid dikaryotic | PRJNA1251115 |

**Table S4.** Number of heterozygous Single Nucleotide Polymorphism (SNPs) found when aligning the HiFi reads to the genome of four basidiomycetes with putative dikaryotic haploid genomes. Phased blocks represent contiguous regions of heterozygous SNPs assigned to the same haplotype, and the phased block N50 indicates the length at which half of the phased sequence is contained in blocks of that size or longer.

| Genomes | Total number of heterozygous SNPs | #Fraction of phased variants | # phased blocks | Phased blocks N50 (Kb) |
| --- | --- | --- | --- | --- |
| *Basidioascus undulatus* | 2,900 | 12% | - | - |
| *Holtermanniella festucosa* | 36 | 75% | - | - |
| *Pholiota brunnescens* | 1,360,152 | 98% | 282 | 657 |
| *Lyophyllum atratum* | 1,072,632 | 99% | 187 | 870 |

**Table S5.** Statistics for the mitochondrial genome assemblies of 18 pyrophilous fungi sequenced using PacBio HiFi technology

| **Species** | **# contigs** | **Total length (Kbp)** |
| --- | --- | --- |
| *Penicillium atrosanguineum* | 1 | 32.7 |
| *Penicillium antarcticum* | 1 | 49.8 |
| *Aspergilus cf. fellis* | 1 | 30 |
| *Rasamsonia columbiensis* | 1 | 47.3 |
| *Peziza echinospora* | 1 | 23.6 |
| *Anthracobia macrocystis* | 1 | 21.8 |
| *Geopyxis carbonaria* | 1 | 141.5 |
| *Pyronema omphalodes* | 1 | 215.8 |
| *Morchella eximia* | 3 | 116.1 |
| *Tricharina praecox* | 3 | 267.4 |
| *Alternaria* sp. | 1 | 123.6 |
| *Coniochaeta hoffmannii* | 1 | 28 |
| *Neurospora sp.* E-DF1 | 2 | 146.4 |
| *Neurospora discreta* E-DF3 | 1 | 71.2 |
| *Holtermanniella festucosa* | 1 | 38.5 |
| *Basidioascus undulatus* | 1 | 81 |
| *Lyophyllum atratum* | 6 | 144.3 |
| *Pholiota brunnescens* | 7 | 197.1 |

**Table S6.** Number of RNA-seq reads generated for gene annotation and resulting gene coding coverage, the number of protein coding genes predicted for each genome, and their BUSCO scores. The times (X) of gene coding coverage by the RNA-seq reads was calculated by dividing the number of RNA-seq bases divided to the whole transcriptome length.

| **Species** | **# Paired-end RNA-seq reads** | **Gene coding coverage (X)** | **# protein-coding genes** | **BUSCO genes % (C: complete (duplicated), F: fragmented, M: missing)** |
| --- | --- | --- | --- | --- |
| *Penicillium atrosanguineum* | 10,598,042 | 100 | 10,889 | C:99.5 (0.5), F:0.1, M:0.4 |
| *Penicillium antarcticum* | 9,428,190 | 80 | 11,884 | C:99.4 (0.5), F:0.2, M:0.4 |
| *Aspergilus cf. fellis* | 8,716,208 | 75 | 12,511 | C:99.7 (0.2), F:0.2, M:0.1 |
| *Rasamsonia columbiensis* | 10,892,180 | 99 | 11,431 | C:98.2 (0.4), F:0.3, M:1.5 |
| *Peziza echinospora* | 9,726,640 | 85 | 12,612 | C:96.4 (0.1), F:0.6, M:3.0 |
| *Anthracobia macrocystis* | 12,948,684 | 102 | 14,887 | C:98.1 (3.5), F:0.5, M:1.4 |
| *Geopyxis carbonaria* | 9,688,470 | 93 | 12,347 | C:90.0 (0.2), F:1.7, M:8.3 |
| *Pyronnema omphalodes* | 9,719,385 | 97 | 10,780 | C:97.2 (1.1), F:0.5, M:2.3 |
| *Morchella eximia* | 11,653,119 | 105 | 11,595 | C:98.9 (1), F:0.2, M:0.9 |
| *Tricharina praecox* | 12,397,547 | 121 | 10,699 | C:97.7 (0.5), F:0.4, M:1.9 |
| *Alternaria* sp. | 13,532,530 | 103 | 13,728 | C:99.0 (0.1), F:0.3, M:0.7 |
| *Coniochaeta hoffmannii* | 15,307,309 | 127 | 12,275 | C:99.7 (0.3), F:0.1, M:0.2 |
| *Neurospora sp.* E-DF1 | 15,349,889 | 147 | 9,749 | C:95.5 (0.2), F:0.2, M:4.3 |
| *Neurospora discreta* E-DF3 | 14,170,355 | 137 | 9,777 | C:99.1 (0.1), F:0.1, M:0.8 |
| *Holtermanniella festucosa* | 2,738,503 | 37 | 7,097 | C:98.0 (0.2), F:0.5, M:1.5 |
| *Basidioascus undulatus* | 13,132,338 | 134 | 11,623 | C:98.2 (0.4), F:0.3, M:1.5 |
| *Lyophyllum atratum* | 9,222,523 | 65 | 14,890 | C:97.5 (7.1), F:0.5, M:2.0 |
| *Pholiota brunnescens* | 13,568,008 | 95 | 16,135 | C:97.3 (1), F:0.7, M:2.0 |

**Table S7.** Test of phylogenetic signature for the enrichment of aromatic C degradation and N acquisition genes in pyrophilous fungi. Using a rooted phylogenetic tree generated by Orthofinder and gene counts normalized by total proteome size of each genome, we quantified phylogenetic signal for each gene with Pagel’s λ, estimating false discovery rate (FDR) adjusted P-value.

| Genes | Gene enrichment  FDR adjusted p-value |
| --- | --- |
| *nahD* | 3.25E-13 |
| *nahB_doxE* | 2.23E-14 |
| *nahA* | 1.89E-17 |
| *nahF* | 4.24E-18 |
| *pcaL_pcaD* | 1.13E-14 |
| *fadA_fadI* | 2.93E-10 |
| *pcaF* | 4.61E-16 |
| *mhpD* | 8.62E-15 |
| *mhpE* | 3.42E-08 |
| *praC_xylH* | 3.29E-17 |
| *mhpF* | 2.88E-21 |
| *catA* | 7.64E-22 |
| *catB* | 1.23E-12 |
| *ligK_galC* | 0.006 |
| *ligA_LigB* | 1.69E-07 |
| *pcaB* | 5.09E-22 |
| *pcaC* | 8.97E-10 |
| *pcaH_pcaG* | 7.70E-09 |
| *nosZ* | 2.59E-16 |
| *nirK* | 6.73E-16 |
| *narG_narZ* | 7.24E-15 |
| *P450nor* | 1.42E-27 |

**Table S8.** GenBank accessions and contig N50 and of 38 public genomes used for comparative genomic analysis

| **Isolate name** | **GenBank accession** | **Contig N50 (Mb)** |
| --- | --- | --- |
| *Alternaria ethzedia* BMP0044 | GCF_023757985.1 | 0.9 |
| *Alternaria incomplexa* BMP0042 | GCF_024043165.1 | 1.7 |
| *Alternaria metachromatica* BMP0045 | GCF_023757995.1 | 1.0 |
| *Ascodesmis nigricans* CBS389.68 | GCA_004786065.1 | 0.1 |
| *Aspergillus felis* CNM-CM5623 | GCA_014281895.1 | 0.1 |
| *Aspergillus fumigatus* Af293 | GCF_000002655.1 | 2.5 |
| *Aspergillus lentulus* IFM54703 | GCA_001445615.2 | 4.2 |
| *Aspergillus pseudoviridinutans* IFM55266 | GCF_018340605.1 | 1.1 |
| *Coniochaeta hoffmannii* EXF-13287 | GCA_030052785.1 | 0.1 |
| *Coniochaeta ligniaria* NRRL30616 | GCA_001879275.1 | 0.5 |
| *Coniochaeta pulveracea* CAB683 | GCA_003635345.1 | 0.3 |
| *Coniochaeta sp*. PMI546 | GCA_021432605.1 | 0.7 |
| *Flammula alnicola* AH47727 | GCA_015499995.1 | 0.2 |
| *Kalaharituber pfeilii* F3 | GCA_015179045.1 | 0.6 |
| *Lyophyllum shimeji* AT787 | GCA_026008595.1 | 1.6 |
| *Morchella conica* CCBAS932 | GCA_003790465.1 | 0.1 |
| *Morchella importuna* M04M26 | GCF_003444635.1 | 1.0 |
| *Neurospora crassa* FGSC 4830 | GCA_033458375.1 | 2.3 |
| *Neurospora hispaniola* FGSC10403 | GCF_033458365.1 | 4.3 |
| *Neurospora tetrasperma* FGSC2508 | GCF_000213175.1 | 0.1 |
| *Neurospora tetraspora* CBS560.94 | GCF_033439725.1 | 4.9 |
| *Penicillium antarcticum* IBT31339 | GCF_028974205.1 | 3.8 |
| *Penicillium atrosanguineum* IBT20685 | GCF_028827265.1 | 3.9 |
| *Penicillium bovifimosum* IBT22155 | GCF_028826915.1 | 3.3 |
| *Penicillium brevicompactum* IBT35673 | GCA_028827255.1 | 6.1 |
| *Penicillium citrinum* IBT23319 | GCF_028827155.1 | 3.8 |
| *Penicillium maclennaniae* IBT15551 | GCF_028827695.1 | 3.6 |
| *Penicillium waksmanii* IBT27052 | GCF_028829765.1 | 4.9 |
| *Pholiota conissans* CIRM-BRFM 674 | GCA_015484465.1 | 0.1 |
| *Pyronema domesticum* CBS144463 | GCA_024516145.1 | 1.5 |
| Rasamsonia emersonii CBS393.64 | GCF_000968595.1 | 0.1 |
| *Sphaerosporella brunnea* GMNB300 | GCA_008704415.1 | 0.1 |
| *Talaromyces atroroseus* IBT11181 | GCF_001907595.1 | 0.4 |
| *Talaromyces proteolyticus* PMI201 | GCF_021365285.1 | 2.6 |
| *Tirmania nivea* G3 | GCA_015179035.1 | 0.6 |
| *Tricholomella constricta* CBS661.87 | GCA_013368375.1 | 0.9 |
| *Trichophaea hybrida* UTF0779 | GCA_015178995.1 | 0.2 |
| *Wilcoxina mikolae* CBS423.85 | GCA_014904905.1 | 0.1 |

**Table S9.** Genes for acquisition of aromatic carbon (C) degradation indicated by † and nitrogen (N), their corresponding pathways, and number of public proteins used for constructing Hidden Markov Model (HMM) for annotating the gene in pyrophilous fungal and their sister species.

| Gene | Description | Pathways | #protein used for creating HMM profile |
| --- | --- | --- | --- |
| *nahD* | 2-hydroxychromene-2-carboxylate-isomerase | Naphthalene_degredation† | 160 |
| *nahB_doxE* | cis-1,2-dihydro-1,2-dihydroxy naphthalene | Naphthalene_degredation† | 12 |
| *nahA* | Naphthalene-1,2-dioxygenase subunit-alpha | Naphthalene_degredation† | 52 |
| *nahF* | Salicylaldehyde_dehydrogenase | Naphthalene_degredation† | 392 |
| *pcaL_pcaD* | 3-oxoadipate-enol-lactonase | bKA_Pathway† | 422 |
| *fadA_fadI* | Acetyl-Co A acyltransferase | bKA_Pathway† | 348 |
| *pcaF* | Beta-ketoadipyl-CoA-thiolase | bKA_Pathway† | 624 |
| *mhpD* | 2-keto-4-pentenoate hydratase | Catechol_Metacleavage† | 191 |
| *mhpE* | Hydroxy_oxovalerate_aldolase | Catechol_Metacleavage† | 81 |
| *praC_xylH* | 4-oxalocrotonate_tautomerase | Catechol_Metacleavage† | 189 |
| *mhpF* | Acetaldehyde_dehydrogenase | Catechol_Metacleavage† | 31 |
| *catA* | Catechol-1,2-dioxygenase | Catechol_Orthocleavage† | 480 |
| *catB* | Muconate cycloisomerase | Catechol_Orthocleavage† | 370 |
| *ligK_galC* | 4-hydroxy-4-methyl-2-oxoglutarate aldolase | PCA_Metacleavage† | 208 |
| *ligA_LigB* | Protocatechuate-dioxygenase | PCA_Metacleavage† | 63 |
| *pcaB* | 3-carboxy-cis,cis-muconate-cycloisomerase | PCA_Orthocleavage† | 18 |
| *pcaC* | 4-carboxy muconolactone-decarboxylase | PCA_Orthocleavage† | 1048 |
| *pcaH_pcaG* | Protocatechuate-3,4-dioxygenase | PCA_Orthocleavage† | 277 |
| *nosZ* | Nitrous-oxide-reductase | N_acquisition | 50 |
| *nirK* | Nitrite-reductase | N_acquisition | 1577 |
| *narG_narZ* | Nitrate-reductase | N_acquisition | 2552 |
| *P450nor* | P450_nor (nitric oxide reductase) | N_acquisition | 263 |

| Genes | Variant enrichment  FDR adjusted p-value |
| --- | --- |
| pcaH_pcaG(S) | 9.46E-20 |
| pcaH_pcaG(N) | 1.12E-19 |
| narG_narZ(S) | 7.70E-18 |
| narG_narZ(N) | 4.69E-15 |
| CatA(S) | 4.69E-15 |
| CatA(N) | 2.63E-19 |
| praC-xyIH(S) | 1.69E-14 |
| praC-xyIH(N) | 5.40E-09 |
| fadA_fadI(S) | 5.31E-10 |
| fadA_fadI(N) | 0.011489 |
| nahD(S) | 8.97E-19 |
| nahD(N) | 9.46E-20 |

Table S10. Test of phylogenetic signature for the genetic variants in aromatic C degradation and N acquisition genes in pyrophilous fungi. Using a rooted phylogenetic tree generated by Orthofinder and gene counts normalized by total proteome size of each genome, we quantified phylogenetic signal for each gene with Pagel’s λ, estimating false discovery rate (FDR) adjusted P-value.
